# Identifying the best approximating model in Bayesian phylogenetics: Bayes factors, cross-validation or wAIC?

**DOI:** 10.1101/2022.04.22.489153

**Authors:** Nicolas Lartillot

**Affiliations:** Laboratoire de Biométrie et Biologie Evolutive, UMR CNRS 5558, Université Lyon 1, Villeurbanne, France

**Keywords:** cross-validation, Bayes factor, marginal likelihood, model comparison, wAIC

## Abstract

There is still no consensus as to how to select models in Bayesian phylogenetics, and more generally in applied Bayesian statistics. Bayes factors are often presented as the method of choice, yet other approaches have been proposed, such as cross-validation or information criteria. Each of these paradigms raises specific computational challenges, but they also differ in their statistical meaning, being motivated by different objectives: either testing hypotheses or finding the best-approximating model. These alternative goals entail different compromises, and as a result, Bayes factors, cross-validation and information criteria may be valid for addressing different questions. Here, the question of Bayesian model selection is revisited, with a focus on the problem of finding the best-approximating model. Several model selection approaches were re-implemented, numerically assessed and compared: Bayes factors, cross-validation (CV), in its different forms (k-fold or leave-one-out), and the widely applicable information criterion (wAIC), which is asymptotically equivalent to leave-one-out cross validation (LOO-CV). Using a combination of analytical results and empirical and simulation analyses, it is shown that Bayes factors are unduly conservative. In contrast, cross-validation represents a more adequate formalism for selecting the model returning the best approximation of the data-generating process and the most accurate estimates of the parameters of interest. Among alternative CV schemes, LOO-CV and its asymptotic equivalent represented by the wAIC, stand out as the best choices, conceptually and computationally, given that both can be simultaneously computed based on standard MCMC runs under the posterior distribution.

## Introduction

Anyone who has worked on data analysis for addressing phylogenetic or evolutionary questions has been faced with the question of selecting among alternative statistical models. For a given problem, one is often faced with several approaches, entailing different assumptions and sometimes returning different estimates for the phylogeny or the quantity of interest. One is then left with the question of how to interpret the differences and which model to favor. The question is difficult for several reasons. First, the models of interest typically differ in their parameterisation, both in structure and in dimensionality, preventing direct comparison of their likelihood scores and requiring careful formalization of how to penalize them accordingly. Second, on a more conceptual front, model selection can be motivated by different objectives, depending on the specific question of interest. These alternative goals entail different compromises and may therefore imply different model selection procedures.

In some cases, the goal of model selection is to test alternative hypotheses about the underlying mechanisms. A relevant example in molecular evolution is the problem of determining whether or not a gene is under positive selection, using phylogenetic codon models. Two alternative models are confronted, one that allows for sites and/or branches to evolve under a positive selection regime, tested against a null model that only allows for purifying selection (Nielsen & Yang, 1998; Zhang *et al*., 2005; Kosakovsky Pond & Frost, 2005). Another example in phylogenetics is the test for the monophyly of a clade. In these examples, the alternative models being considered are meant to be idealized representations of alternative possible states of nature. As a result, the aim is to identify the ‘true’ model, i.e. the model formally representing the true objective situation.

In a classical frequentist context, the standard approach to deal with such hypothesis testing problems is to use likelihood ratio tests, relying on chi-square asymptotics or on parametric (Goldman, 1993) and non-parametric (Shimodaira, 2004) approaches to approximate the distribution under the null. In a Bayesian context, hypothesis testing can be addressed in two different ways. One approach is to compare the marginal likelihoods under the two models, or equivalently, to compute the Bayes factor, i.e. the ratio of the two marginal likelihoods (Jeffreys, 1935; Kass & Raftery, 1995; Oaks *et al*., 2019). Alternatively, a fully Bayesian formalization of the problem suggests to also define a prior probability over the models and then to select models based on their posterior probabilities (Kass & Raftery, 1995).

In other situations, the question is instead to select the model that gives the most accurate estimation or the best approximation for the data-generating process, and this, without consideration of any hypothesis that would be true or false. A paradigmatic example is to choose the degree of a polynomial regression function (see e.g. Burnham & Anderson, 2002). Here, the true regression function is not generally believed to be itself a polynomial, and thus there is no question of identifying the true degree. Instead, the question is to find the best tradeoff between the lack of flexibility of polynomials of lower degree and the increased estimation error entailed by a higher degree.

In phylogenetics, instances of this second version of model selection are often encountered. An example is the problem of choosing between an empirical model of amino-acid evolution such as JTT (Jones *et al*., 1992), WAG (Whelan & Goldman, 2001) or LG (Le & Gascuel, 2008), or the general time reversible (GTR) model. Empirical matrices are estimates of the average amino-acid exchange rates across a heterogeneous set of proteins and taxonomic groups. As a result, the biochemical prior information that they encode will fit a specific dataset of interest only approximately. If the dataset of interest happens to be sufficiently large, re-estimation of the complete general time-reversible model may give a more accurate model than the one proposed by any available empirical matrix. The problem that model selection has to solve in this context is whether one can afford this parameterization or whether falling back onto the prior biochemical knowledge encoded into an empirical matrix represents a safer option. The answer to this question will fundamentally depend on data size, but also, on how well the biochemical information encoded into currently available amino-acid replacement matrices generalizes to the specific dataset of interest.

As another example, accounting for pattern heterogeneity across sites is usually done using mixture models (Pagel & Meade, 2004; Lartillot & Philippe, 2004; Evans & Sullivan, 2012; Susko *et al*., 2018; Schrempf *et al*., 2020). In that context, the question of model selection is important, and non trivial, whether for choosing between alternative empirical models, for determining the number of components, or for the sake of a more general assessment of alternative mixture designs. However, the true distribution of substitution rates or patterns across sites is not itself a mixture. Instead, the hope is just that a well chosen mixture should give a reasonable approximation of the unknown true distribution, which would then provide increased robustness for phylogenetic inference purposes. The situation is thus formally similar to the one described above in a regression context using a polynomial regression function: the point of model selection with these phylogenetic mixture models is not to identify the true number of components, but to find a good compromise between the lack of flexibility of model with few mixture components, and the increased estimation error incurred under rich mixtures.

The general problem of finding the best approximating model, as opposed to testing hypotheses, has been classically formalized in different ways. On one side, the approaches used for hypothesis testing, namely likelihood ratio tests and Bayes factors, have often been employed in this context as well. However, it is not totally clear whether they represent a correct formalization of the question, given that there is no proper hypothesis to be tested. As pointed out by Akaike (1974) and others (Burnham & Anderson, 2002; Sullivan & Joyce, 2005), hypothesis testing is not adequately formulated, in decision-theoretic terms, as a procedure of approximation, the two goals being intrinsically different. In the more specific context of Bayesian inference, Bayes factors or model posterior probabilities have been recognized as appropriate only in circumstances where it was believed that one of the competing models was in fact true, and that in other circumstances, other criteria may be more appropriate (Bernardo & Smith, 1994; Konishi & Kitagawa, 2007). Accordingly, approaches have been developed, which are more decisively framing the question in terms of finding the best approximation, without predicating on any model being true. Among these approaches, two main types can be identified: cross-validation and information criteria.

The idea of cross-validation is to train the model on a subset of the data and then evaluate the fit of the model over another non-overlapping subset of the observations. The procedure is typically repeated over multiple random splits of the data into a training and a validation set, and the score is averaged over these replicates (Stone, 1974; Zhang, 1993; Smyth, 2000; Geisser, 1975; Geisser & Eddy, 1979; Gelfand, 1996). Given its operational definition, cross-validation thus directly estimates the predictive fit of a model. However, this apparent focus on the predictive performance does not imply that cross validation will be useful only in a context where prediction is indeed contemplated in practice. Perhaps a more fundamental justification is the following: since good prediction of future data can be achieved only by capturing, through the fine-tuning of the parameters of the model, the structural features of the data-generating process, the predictive fit should be a good indicator of estimation accuracy. By a similar argument, it can also be seen that cross-validation automatically accounts for overfitting. By definition, overfitting is what happens when a model captures random, non-reproducible patterns in the data. Owing to this non-reproducibility, a model that overfits will therefore show a poor fit on new data obtained from the same population. This idea can be quantitatively formalized in terms of the generalization gap of a model (Thomas *et al*., 2020), or optimism (Efron, 1986), which is defined as the average drop in the apparent log-likelihood score, when going from the training set to the validation set. Altogether, more complex models will thus have more expressiveness for capturing structural features of the data-generating process, but they will also tend to have a wider generalization gap. Cross-validation automatically captures the balance between these two opposing components of the overall fit.

In the details, cross-validation can be implemented in many different ways, depending on what proportion of the data to set aside for validation, or how many replicates to consider (see Zhang, 1993, for an overview). The simplest and original approach is leave-one-out cross-validation (LOO-CV), whereby each observation is successively taken out of the sample and reserved for subsequent validation of the model, while training is done on the remaining data (Stone, 1974). Alternatively, in k-fold cross validation (k-fold CV), the dataset is split into *k* equal sized subsets, then each subset is set aside for validation and the remaining *k* − 1 subsets are used for training (Breiman *et al*., 1984; Zhang, 1993).

In all cases, direct implementation of cross-validation is expensive, owing to the total number of replicates to consider. A variant of k-fold cross-validation has previously been used in a phylogenetic context (Lartillot *et al*., 2007; Lartillot & Philippe, 2008), sometimes in combination with strict subsampling, i.e. using training and validation sets that together represent a subset of the data (Pisani *et al*., 2015). Strict subsampling was motivated by the need to reduce the computational cost. A downside, however, is that the models are then under a regime of data size that does not correspond to the effective regime in which subsequent inference is conducted. Yet the relative fit of alternative models with differing dimensions depends on data size, since higher-dimensional models typically require more data to learn their parameters.

For all these reasons, indirect approaches to cross-validation, which would avoid the explicit resampling and fitting procedure, would be particularly useful. In this direction, and in the specific case of leave-one-out, it is in fact possible to get an estimate of the cross-validation score based only on a standard MCMC run conditioned on the full dataset (Gelfand *et al*., 1992; Chen *et al*., 2012; Lewis *et al*., 2014). This clever importance sampling approach, called cross-predictive ordinates (CPO), makes leave-one-out cross-validation particularly attractive in a Bayesian context.

In a more theoretical spirit, and starting with Akaike (1974), a long series of information criteria have been proposed, based on information-theoretic considerations. The fundamental idea behind these information criteria is to identify the model which, once trained on the dataset of interest, induces a distribution over the data that is closest to the true distribution of the population. Mathematically, the distance between model and truth is measured by the information loss (i.e. the Kullback-Leibler divergence). Importantly, this distance is measured under the effective conditions of use of the model, that is, under the current data size. As a result, it accounts for the two different reasons why the model might not be so close to the true distribution in practice: because of model mis-specification, but also, because of stochastic error in parameter estimation due to finite sample size. This last point will critically depend on both the size of the dataset and the model dimension.

Information criteria were first derived in a maximum likelihood context. The original criterion proposed by Akaike, the AIC, has a particularly simple expression. However, its derivation relies on the assumption that the models being considered are not far from the true distribution. It is thus not valid under strong model violation, a situation often encountered in practice. The AIC was revisited by Takeuchi (in an original contribution in Japanese, as reported in Konishi & Kitagawa, 1996), who proposed a criterion, the TIC, which is valid even in the presence of strong model violation. The TIC reduces to the AIC when the data are indeed under the model for some true parameter value. Compared to the AIC, the TIC is slightly more involved computationally. In practice, however, the difference between TIC and AIC can be substantial (Konishi & Kitagawa, 1996).

The TIC was then adapted to the maximum penalized likelihood framework, with the regularized information criterion (RIC, Shibata, 1989). A more general derivation, valid for a broader range of plug-in estimators (including maximum likelihood, maximum penalized likelihood or posterior mean Bayesian point estimate), was proposed, in the form of the generalized information criterion (GIC, Konishi & Kitagawa, 1996). Finally, the widely applicable information criterion (wAIC Watanabe, 2009) represents a more specific Bayesian adaptation, which, unlike the GIC, does not rely on a plug-in estimator. All of these information criteria, RIC, GIC, and wAIC, provide a measure of the expected predictive fit under the corresponding estimation method, all of which are valid even under model violation, thus like the TIC (and unlike the AIC). The wAIC is also valid under a broader class of models, such as mixture models or Bayesian networks, which are singular, in the sense that they entail some redundancy (i.e. non-identifiability) in the mapping from parameters to probability distributions over the data (Watanabe, 2007). Because of their non-identifiability, such singular models typically have complex asymptotic properties that are not correctly handled by current information criteria. Addressing these complications is what led to the development of singular statistical learning theory (Watanabe, 2001, 2009), of which the wAIC is one of the specific contributions.

Several other information criteria have been proposed, in addition to those mentioned above. Two of them were explicitly meant for Bayesian inference: the deviance information criterion, or DIC (Spiegelhalter *et al*., 2002) and the Bayesian analogue of AIC, or AICM (Raftery *et al*., 2007; Gelman *et al*., 2014). The DIC has been somewhat controversial (Plummer, 2008; Spiegelhalter *et al*., 2014; Celeux *et al*., 2006; Gelman *et al*., 2014). One problem is that it relies on the posterior mean point estimate, which is not easily defined for mixture models (Celeux *et al*., 2006) or in a phylogenetic context. Another problem is that, like the AIC, the DIC assumes that the model is correctly specified (Spiegelhalter *et al*., 2002). As for the AICM, it was derived based on an analogy with the AIC, by relying on a definition of the effective number of parameters of a model based on the Monte Carlo variance of the log likelihood. There are two problems with this derivation, however. First, the analogy with the AIC, which is a maximum likelihood criterion, fails to capture the contribution of the prior to the fit of a model in the Bayesian case. Second, just like the AIC and the DIC, the AICM does not account for the impact of model violation. Finally, the Bayesian information criterion, or BIC (Schwarz, 2006) represents one last criterion, which does not proceed from the same rationale as the other criteria mentioned above, as it is not based on an information loss argument. Instead, it is meant as an asymptotic expression for the log of the marginal likelihood. As such it is more appropriate for true model identification than for best model approximation purposes (Aho *et al*., 2014). Of note, the BIC can be strongly conservative even in a true model identification task (Vrieze, 2012).

There is a intimate connection between information criteria based on the information loss (AIC, TIC, RIC, GIC, wAIC) and cross-validation. Information criteria are effectively asymptotic approximations of the expected predictive fit of the model trained on a dataset of the original size. Cross-validation, on the other hand, is a direct operational estimate of the expected predictive fit of the model trained on a subset of the data. In the case of LOO-CV, for large datasets, this difference is relatively minor, as leaving out one single data point will have a minor impact on the training. As a result, LOO-CV is asymptotically equivalent to information criteria of the Akaike family (Stone, 1977; Watanabe, 2010*a*), or equivalently, information criteria of the AIC family are just asymptotic expressions for the cross-validation score, each valid under different specific assumptions. This result is important, as it emphasizes the operational meaning of information criteria. Practically, it suggests simple experiments on empirical data, to check the range of data size under which this asymptotic equivalence is effective.

Altogether, there is thus by now a broad theoretical background on model selection. Several alternative methods have been proposed, with subtle differences concerning their aim or their exact regime of applicability. These issues have already been discussed in the applied statistical literature (Burnham & Anderson, 2002; Aho *et al*., 2014; Vrieze, 2012; Konishi & Kitagawa, 2007), yet this has not yet been fully incorporated into current phylogenetic practice. This is particularly apparent in Bayesian phylogenetics. Thus, although it has long been noted that Bayes factors are conservative in model selection when used in combination with vague priors on the model specific parameters (the so-called Jeffreys-Lindley paradox, Jeffreys, 1967; Lindley, 1957), and that cross-validation approaches may be more adequate for best-approximating model selection (Gelfand *et al*., 1992; Bernardo & Smith, 1994; Konishi & Kitagawa, 2007), Bayes factors or marginal likelihoods are often presented as the method of choice (Kass & Raftery, 1995; Lartillot & Philippe, 2006; Xie *et al*., 2011; Oaks *et al*., 2019) and are widely used (Suchard *et al*., 2001; Baele *et al*., 2012*b*, 2013; Baele & Lemey, 2013; Brown & Thomson, 2017; Ronquist *et al*., 2021). The computational challenges raised by the numerical evaluation of marginal likelihoods (Lartillot & Philippe, 2006; Xie *et al*., 2011; Baele *et al*., 2012*a*) have also represented a clear limitation, which has prevented a broader and more systematic application of this paradigm to current empirical problems based on large datasets. Cross-validation was used in Bayesian phylogenetics primarily for computational reasons (Lartillot *et al*., 2007; Lartillot & Philippe, 2008), although without any correct evaluation of its numerical accuracy and its theoretical validity in that context. The implementation of LOO-CV offered by CPO appears to be attractive, and has already been introduced specifically in phylogenetics (Lewis *et al*., 2014), but has thus far not been broadly used in this context. Finally, the wAIC has never been applied to phylogenetic model selection.

In this work, the theoretical and methodological background just presented is utilized to revisit the question of Bayesian model selection in phylogenetics, with a focus on the question of choosing among alternative models of sequence evolution, and with an emphasis on identifying the best approximating model, irrespective of any question about hypothesis testing. The statistical and numerical issues are both examined. On the numerical side, the work presented here starts from the realization that k-fold cross-validation, such as implemented in PhyloBayes (Lartillot *et al*., 2013), turns out to be numerically inaccurate. This point is examined, and an alternative method is proposed, based on sequential importance sampling (sIS), which is similar to sequential Monte Carlo (Wang *et al*., 2016) and gives an estimate of the marginal likelihood and, simultaneously, the k-fold cross validation scores for any *k*. This sIS approach is computationally intensive but can be used on datasets of relatively small size to validate and compare marginal likelihood and cross-validation for their ability to select the model that is most accurate in parameter estimation. Finally, the CPO approach to leave-one-out cross-validation is re-implemented, its statistical and numerical properties are characterized, and its connection with the wAIC is explored on an empirical phylogenomic dataset.

## Materials and Methods

### Definitions and relations between alternative Bayesian measures of model fit

In this subsection, the alternative Bayesian measures of model fit are formally defined. A homogeneous mathematical notation is introduced, so as to emphasize the connections and the differences between them, leaving aside in a first step the numerical and algorithmic problems.

Suppose we have a dataset made of *n* observations, *X* = (*X_i_*)_*i*=1..*n*_. In the context of phylogenetic inference, these observations would typically be the columns of a multiple sequence alignment. In the following, we will adopt a frequentist perspective and assume that these observations are iid from an infinite population of unknown distribution.

We then consider a model *M* parameterized by *θ*. In a Bayesian framework, this model is endowed with a prior *p*(*θ*) and then conditioned on data *X*, giving the posterior distribution *p*(*θ* | *X*):

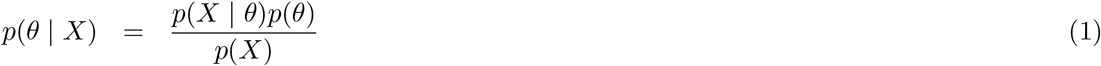

where

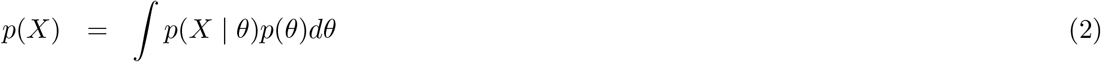

is the marginal likelihood. We wish to evaluate the fit of the model.

#### Marginal likelihoods and the Bayes factor

A first approach is to use the marginal likelihood as the measure of the fit. In the following, when multiple models are compared, the dependence of the marginal likelihood on the specific model will be more explicitly noted *p*(*X* | *M*), for model *M*. Otherwise, the simpler notation *p*(*X*) is used. Often, the fit of a given model *M*_2_ is computed relatively to another model *M*_1_, by computing the Bayes factor, defined as the ratio of the marginal likelihoods of the two models (Jeffreys, 1935):

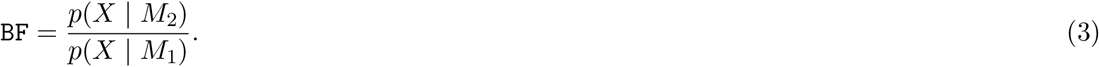

A Bayes factor greater than 1 thus means that model *M*_2_ has a higher fit, compared to *M*_1_. As a way to ensure a scaling consistency across all alternative measures of fit considered here, it is useful to define the per-site log marginal likelihood:

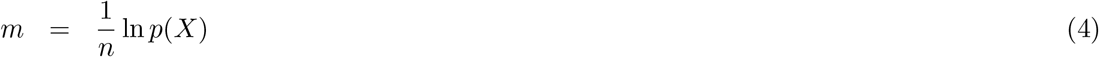

or, when comparing two models, the per-site log Bayes factor:

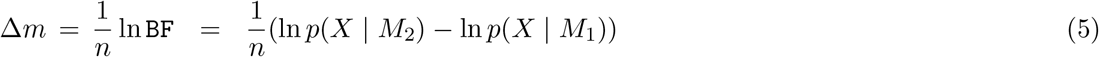

#### Bayesian cross-validation

An alternative to marginal likelihoods and Bayes factors is cross-validation. As mentioned in the introduction, the general idea is to split the dataset into two subsets, using one subset (noted *X^t^*) for training the model and then evaluating the fit of the model over the remaining subset (noted *X^v^*, for validation). In the context of Bayesian inference, a natural procedure to implement cross-validation is to average the likelihood under the validation set over the posterior distribution under the training set. The resulting cross-validation score is then log-transformed and averaged over multiple random splits of the original dataset into training and a validation sets. Of note, other approaches have been proposed, such as computing the cross-validated likelihood on a plugin estimate of the model’s parameters, typically, the posterior mean (Konishi & Kitagawa, 1996). However, this approach is not applicable for singular and redundant models, such as mixture models (Plummer, 2008).

Based on this general idea, multiple settings can be contemplated for implementing cross-validation. These alternative settings differ in how the dataset is split, how the replication procedure is defined, or whether the likelihood is averaged over the posterior distribution jointly for all observations of the validation set, or independently for each of them.

In *k*-fold cross-validation, the dataset is split into *k* subsets of equal size. Then, each subset is considered in turn as the validation set, while the other *k* − 1 subsets are pooled together to make the training set. There are thus *k* replicates in total. In a variant of this approach, previously used in phylogenetics (Lartillot & Philippe, 2008), each replicate is obtained independently of other replicates, by randomly splitting the dataset into a fraction *f* of the observations, which is set aside for validation, while the remaining fraction 1 − *f* used for training. It is thus close to the original version of *k*-fold cross-validation, with *f* = 1/*k*, except that the replicates are not obtained by systematic rotation of the subsets. As a result, the number of replicates can be arbitrary. In practice, for computational reasons, a small number of replicates is used, typically *m* = 10. The fraction *f* is typically set to 0.1 or 0.2, or equivalently, *k* = 10 or 5. In the following, this approach will also be called k-fold cross-validation, even if it does not exactly correspond to the original version.

To more formally describe the detailed procedures, assume that *l* = 1..*L* replicates are considered, each based on a random split of the dataset into 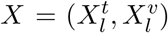, and that the training and validation sets are of size *q* and *r*, respectively. Thus, in k-fold CV, *q* = (1 − *f*)*n* and *r* = *fn*, with *f* = 1/*k*, but these definitions are more generally valid for other cross-validation schemes.

Using these notations, a first version of k-fold cross-validation score, which in the following will be called *joint* k-fold CV, consists in averaging the joint likelihood of all data points of the validation set over the posterior under the training set, i.e. computing, for replicate *l*:

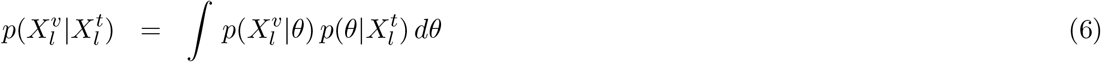

and then averaging the logarithm of this score over the replicates:

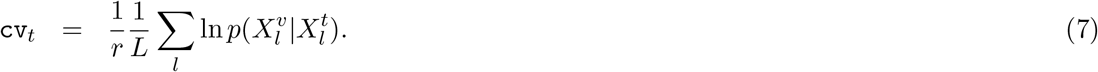

The subscript in cv_*j*_ refers to the fact that the score is based on the joint likelihood of the validation set. Note also that, in this definition, the logarithmic score is divided by the size of the validation set. It is thus a measure of the predictive score per future observation. This convention will be useful for comparing the alternative settings introduced below, which differ in the value of *r*.

The definition just given corresponds to how cross-validation was implemented previously in a phylogenetic context (Lartillot *et al*., 2007; Lartillot & Philippe, 2008). Alternatively, the posterior averaging can be done independently for each observation of the validation set. Noting 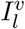 the subset of 1..*n* corresponding to the indices of the observations assigned to the validation set in replicate *l*, the *sitewise* cross-validation score is thus defined as:

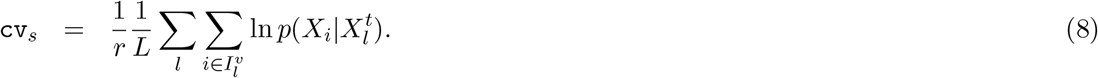

Again, the score is per future observation. The sitewise approach has recently been used in the context of phylogenetics (Bujaki & Rodrigue, 2022).

Finally, in *leave-one-out* cross-validation, each observation is taken in turn and set aside for validation, using the *n* − 1 remaining observations to train the model. Noting *X*(*i*) the training set (of size *n* − 1) obtained by removing observation *i*, the score is defined as:

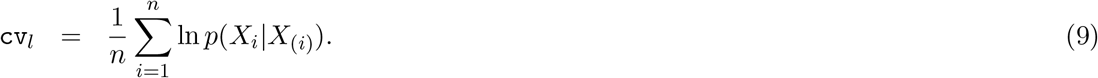

#### The widely applicable (or Watanabe-Akaike) information criterion (wAIC)

As mentioned in the introduction, the idea behind information criteria is to assess models based on how close their predictive distribution (once trained on a current data set) is to the true distribution of the data, using the Kullback-Leibler divergence as a quantitative measure of the deviation between model and truth.

More formally, let *q*\* stand for the unknown true distribution. Then, the Bayes generalization error is defined as the expected Kullback-Leibler divergence between the true distribution and the posterior distribution under the model (Watanabe, 2010*b*):

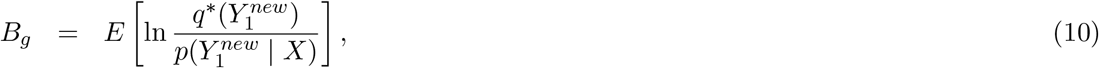

where 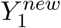 is a single new data point sampled from the population, *X* is a (training) data set of size *n*, and the expectation is over both *X* and 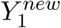. Developing the logarithm gives:

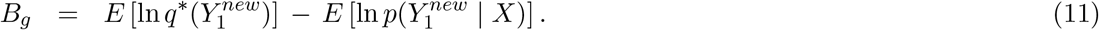

The first term of this equation is the entropy of the empirical source. It does not depend on the model under consideration, and thus, choosing the model minimizing the generalization error *B_g_* is equivalent to choosing the model maximizing the second term, 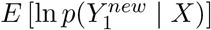, which is just the Bayes predictive fit, i.e. the expected fit of the model on a new data point, once trained on a dataset of size *n*.

The next step of the derivation of information criteria is to derive a practically usable asymptotic approximation of the predictive fit. In this direction, an asymptotic was derived by Watanabe (2007), leading to the wAIC, which takes the following form:

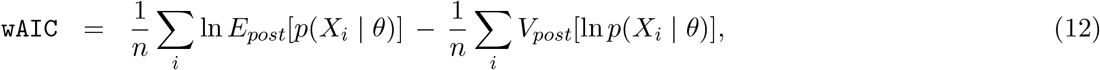

where *E_post_* is the expectation, and *V_post_* the variance, over the posterior distribution under the complete dataset *X*. Of note, the convention used here differs from the classical convention set up by Akaike by a factor −2*n*. That is, the wAIC such as defined here is per site, and higher values corrrespond to a better prediction.

In terms of interpretation, the first term of `wAIC`, which is the average per-site posterior mean likelihood, can be seen as the self-fit, that is, the mean fit of the individual data points of the training set, under the parameter value estimated on that training set. Because it uses the data twice, this measure of the fit is optimistic. The second term, which is the per-site posterior mean variance of the log-likelihood, represents an estimate of this optimism bias. As such, it plays the same role as the dimensional penalty in AIC. Of note, in spite of their similar form, the two terms in equation 12 are not of the same order of magnitude. Owing to the asymptotic concentration of the posterior, the variance of the log likelihood at a typical site (the second term) decreases as a function of data size, whereas the mean log likelihood (the first term) remains asymptotically macroscopic. The same situation holds for the AIC and other classical information criteria, for which the dimensional penalty becomes negligibly small compared to the log likelihood term for sufficiently large datasets.

#### The theoretical targets of Bayesian cross-validation and the wAIC

In this subsection, the theoretical relations between cross-validation (in its different versions) and the wAIC are clarified. Whichever setting is used, from a frequentist perspective, the primary quantity of interest takes the form of an expected predictive fit, which only varies among settings in the size of the training and validation sets. To formalize this point in full generality, define:

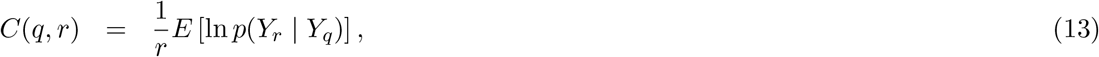

where *Y_q_* and *Y_r_* are two independent datasets randomly drawn from the population, of size *q* and *r*, respectively. In words, *C*(*q, r*) is the expected predictive fit on a dataset of size *r* upon training on an independent dataset of size *q*. By this definition, k-fold joint cross-validation can be seen as an estimator of *C*((1 − *f*)*n, fn*), with *f* = 1/*k*, k-fold site-wise cross-validation as an estimator of *C*((1 − *f*)*n*, 1), leave-one-out cross-validation as an estimator of *C*(*n* − 1, 1), and the wAIC is an estimator of *C*(*n*, 1). The per-site log marginal likelihood is also a special case of this formula, obtained by training on an empty set and validating on the complete dataset, that is, as an estimator of *C*(0, *n*).

Several points deserve discussion here. First, in all cases, the frequentist target *C*(*q, r*), regardless of the specific values of *q* and *r*, is an expectation over data sets randomly sampled from the empirical source. In contrast, all estimators are in practice computed on the unique data set of size *n* that is assumed to be available. However, for large *n*, they all become very close to their respective frequentist targets.

Second, the Bayes generalization error can be written more compactly as:

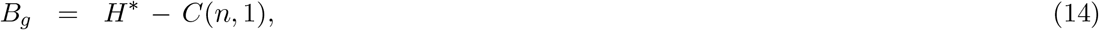

where *H*\* is the entropy of the empirical source. Arguably, minimizing the Bayes generalization error, or equivalently maximizing the Bayes predictive fit *C*(*n*, 1), should be considered as the most meaningful target of model selection in the present context, since in practice, what one wants to evaluate is the fit of the model under the effective regime in which this model is used (i.e. on a dataset of size *n*). Among all explicit cross-validation schemes, LOO-CV is the one closest to this requirement, since it is an estimator of *C*(*n* − 1, 1), which differs from *C*(*n*, 1) by an amount of the order of 1/*n*. For reasonably large datasize, this difference will be negligible. Which of LOO-CV and wAIC is best at approximating the Bayes generalization error is more subtle question. The wAIC is directly targeting *C*(*n*, 1). However, it entails additional approximations, which should vanish asymptotically but could lead to departures for data sets of intermediate size that are bigger than the difference between *C*(*n*, 1) and *C*(*n* − 1, 1). As for k-fold CV, it stands a bit further from the ideal target. In practice, it will tend to underestimate the predictive fit of all models, but more so for those models that are more parameter-rich. To what extent this can make a difference in practice will be explored on several examples below. For more details and more insights about these conceptual issues, and in particular about the difference between joint and sitewise k-fold CV, see also Supplementary Material, section 4.

Finally, if the model is regular, then for large *n*, the posterior distribution becomes increasingly concentrated around an asymptotic parameter value *θ*_0_. In the specific case where the data have been produced under the model, then *θ*_0_ will be the true parameter value. In the general case where the data are from an unknown distribution, there is no true parameter value, in which case *θ*_0_ is the best approximation (in the Kullback-Leibler metric) that the model can give for the distribution induced by this empirical source. In both cases, for large *n*, all expected scores introduced above, k-fold, leave-one-out, marginal or wAIC, converge asymptotically to the expected log likelihood of a single new observation 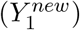 sampled from the population under *θ*_0_:

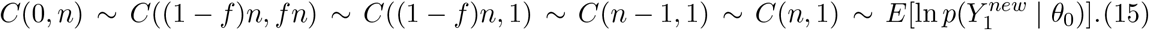

### Monte Carlo methods

To ease notation, in the following, it is assumed that, for cross validation, the first *q* data points were used for training, and the last *r* for validation, with *q* and *r* depending on the exact cross-validation approach. Estimation of the marginal likelihood can be seen as a special case obtained by setting *q* = 0. In what follows, *X_a_*_:*b*_ denotes the set of observations (*X_i_*)_*a*≤*i*≤*b*_. Thus, in particular, *X*_1:*q*_ represents the first *q* observations (i.e. the training set). When *q* = 0, *X*_1:*q*_ is the empty set. The index *i* = 1..*n* runs over data points, and *t* = 1..*T* over the parameter configurations sampled by MCMC.

#### Naive importance sampling (nIS) for k-fold CV

The naive importance sampling (nIS) approach is used for joint and sitewise k-fold CV. In both cases, we assume that an MCMC chain has been run under the training set, yielding a sample (*θ_t_*)_*t*=1..*T*_ approximately under the posterior distribution *θ_t_* ~ *p*(*θ*|*X*_1:*q*_), for *t* = 1..*T*.

First considering joint k-fold cross-validation, equation 6, being an expectation over the posterior distribution under the training set, can be approximated by the corresponding Monte Carlo average:

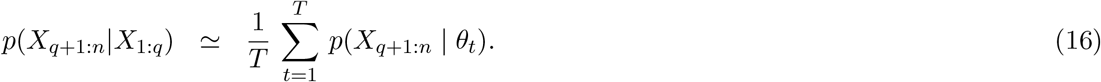

Thus, nIS for joint k-fold CV runs as follows: for *t* = 1..*T*, compute the likelihood of the validation data, *L_t_* = *p*(*X_q_*_+1:*n*_ | *θ_t_*), compute the arithmetic mean of the *L_t_*’s over the *T* Monte Carlo samples and log-transform.

A similar approach can be used for the site-wise version of k-fold CV, since, for any single observation *X_i_* of the validation set:

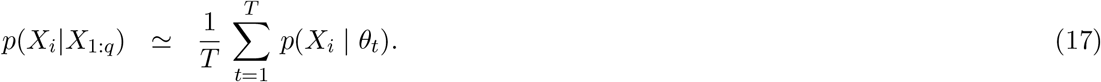

The site-wise posterior averages can be computed in parallel for each observation and then combined according to equation 8. That is, for *t* = 1..*T*, compute the likelihood separately for each data point of the validation data, *L_it_* = *p*(*X_i_* | *θ_t_*), for *i* = *q* + 1..*n*. In a second step, for all *i* = *q* + 1..*n*, compute the arithmetic mean of the *L_it_*’s over the *T* Monte Carlo samples, log-transform, and finally, sum all individual contributions across the *r* data points of the validation set.

#### The cross-predictive ordinate (CPO) approach for LOO-CV

The cross-predictive-ordinate (CPO) approach (Chen *et al*., 2012; Lewis *et al*., 2014) gives an estimate of the leave-one-out cross validation score. It relies on the following harmonic-mean identity:

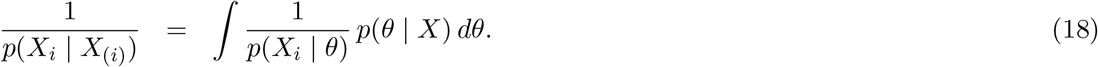

This identity suggests to obtain a sample of parameter configurations from the posterior distribution under the entire dataset, *θ_t_* ~ *p*(*θ*|*X*_1:*n*_), for *t* = 1..*T*, and then approximate the expectation given by equation 18 by a Monte Carlo average:

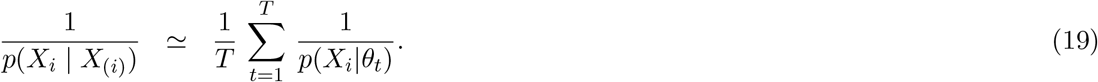

Here also, like for site-wise k-fold CV, the Monte Carlo averages across all sites can be computed in parallel, over a single scan of the MCMC chain. Thus, for each *t* = 1..*T*; for each *i* = 1..*n*, compute the likelihood of each data point separately, *L_it_* = *p*(*X_i_* | *θ_t_*) for site *i*. Then, in a second step, for each site, compute the harmonic mean 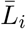 of the *L_it_*’s over the *T* Monte Carlo samples, log-transform and sum all individual contributions across the *r* data points of the validation set.

As a harmonic mean estimator, CPO may be subject to numerical instabilities, caused by the occasional but disproportionate contribution of single MCMC samples for which the site-specific likelihood turns out to be exceptionally small (and thus its inverse particularly large, in equation 19). This is conceptually the same issue as for the harmonic mean estimator of the marginal likelihood (Raftery *et al*., 2007), although the problem is expected to be less dramatic in the present case, where the harmonic mean is applied to the likelihood values at a single site, over parameter configurations sampled from the joint posterior distribution induced by all sites. Still, instabilities may occur, in particular for small data size or for more complex models. To address this point, a stabilized version has been proposed (Vehtari *et al*., 2016), which fits, for each *i*, a generalized Pareto distribution to the right tail of the empirical series of inverse likelihood values (20% largest 1/*L_ik_*’s over the *T* Monte Carlo samples). The contribution of these 20% importance weights to the harmonic mean is then replaced by the expectation under the generalized Pareto distribution. As an aside, the parameter of the fitted Pareto distribution, here noted *κ* (noted *k* in Vehtari *et al*., 2016), offers an additional quality check. According to Vehtari *et al*. (2016), if the value of 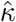 exceeds 0.7, indicating a heavy tail for the inverse likelihood values, then there should be some concern about the reliability of the resulting estimate for the corresponding site. This alternative approach based on Pareto smoothing was not used systematically in the present case, although it was examined more specifically when checking the numerical accuracy of LOO-CV (Results, section Asymptotics of LOO-CV and the wAIC, Supplementary Material, section 3).

#### Sequential Importance Sampling (sIS) for BF and k-fold CV

Sequential Importance Sampling is a step-by-step version of IS, which can be recruited for computing the marginal likelihood of the models of interest, but also, with some minor adaptation, the k-fold CV score for any *k*. It is based on the observation that the joint probability of all data points can be expressed in terms of a sequential product of the marginal likelihoods of each of individual observations:

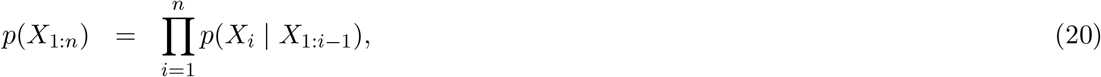

or, on a logarithmic scale and on a per-site basis:

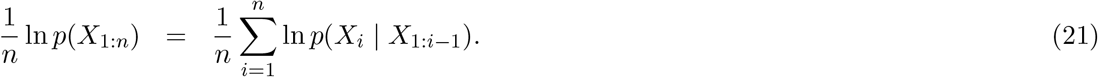

In turn, if, for *t* = 1..*T*, *θ_it_* is sampled from the partial posterior based on the first *i* − 1 data points, i.e. *θ_it_* ~ *p*(*θ* | *X*_1:*i*−1_), then an importance sampling estimate of *p*(*X_i_* | *X*_1:*i*−1_) is given by:

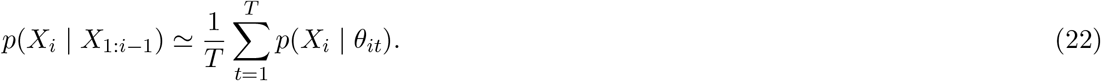

This suggests to run a quasi-static MCMC in which data points are added sequentially, each time running the MCMC for a few cycles and averaging the likelihood of the next data point under parameter configurations sampled from the posterior induced by all current data points (equation 22). These individual IS estimates can then be log-transformed and combined additively (equation 21). This can be seen as a particular case of the stepping-stone approach (Fan *et al*., 2011), in which the interpolation between the prior and the posterior is implemented with partial data sets of increasing size, rather than using power posteriors.

As just mentioned, applying the sIS approach over the complete range of data points gives an estimate of the marginal likelihood. Alternatively, since:

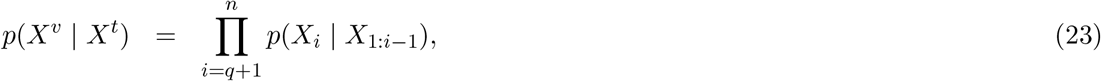

summing only the last *r* data points, with *r* = 1/*k*, gives the joint k-fold CV score. Of note, the order of the data points does not matter for the full marginal likelihood, but it does for joint k-fold CV. In practice, for joint k-fold CV, the procedure is repeated over a series of *L* random permutations of the original dataset, which then automatically implements the *L* replicates, each consisting of a random partition of the original dataset into a training (*i* = 1..*q*) and a validation (*i* = *q* + 1..*n*) sets.

To formalize the practical details of the Monte Carlo procedure for sIS, in the following, a cycle is defined as a coordinated series of multiple MCMC moves that are applied successively on all parameter components of the models. A parameter configuration is saved after each cycle. A cycle can be arbitrary, although in practice, for sIS to give accurate estimates, a cycle should be sufficiently long to give a reasonably good de-correlation of the MCMC between successive saved samples. The algorithm then proceeds as follows. Starting from a parameter configuration sampled from the prior *θ*_0_ ~ *p*(*θ*), at step *i* = 1..*n*:

- the MCMC is run for a short burn-in period of *B* cycles, so as to equilibrate the MCMC, and then for another series of *T* cycles, giving *T* new parameter configurations *θ_it_* approximately under the partial posterior distribution *p*(*θ* | *X*_1:*i*−1_);
- the likelihood of the next data point is calculated under each of these sampled parameter values, i.e. *L_it_* = *p*(*X_i_* | *θ_it_*)
- the arithmetic mean of the *T* likelihood factors *L_it_*, *t* = 1..*T*, is calculated:

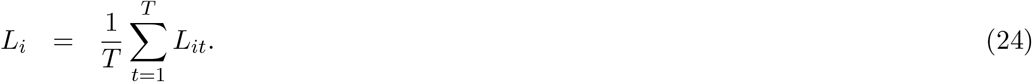

Finally, the *L_i_*’s for *i* = 1..*n* are log-transformed and combined such as specified by equations 21.

In practice, the Monte Carlo procedure just described is not computationally efficient, and this, for two reasons. First, the quality of the estimate of *p*(*X_i_* | *X*_1:*i*−1_) given by equation 24 depends on the variance of the log-likelihood ln *p*(*X_i_* | *θ*) under the partial posterior *p*(*θ* | *X*_1:*i*−1_). When this variance is large, a larger number of samples should be used. In practice, many data points (e.g. constant sites in a phylogenetic context) are characterized by a small variance, while a minority of data points induce a large variance. This suggests that the number of Monte Carlo samples can be tuned on a per-site basis, by first running a small number of cycles at step *i* to estimate the variance of the log-likelihood and then proceed with a number of cycles determined based on this variance estimate (see Supplementary Material, section 1.2).

Second, the estimate of the marginal likelihood given by this approach can have a large variance, which is mostly contributed by the first steps of the algorithm, i.e. when sampling from a distribution close to the prior. A similar problem was encountered by Fan *et al*. (2011). The solution that was then proposed to reduce the variance is to interpolate, not between the prior and the posterior, but between an empirical estimate of the posterior and the true posterior. A similar approach is adopted here. Specifically, the version of the sIS algorithm described thus far is such that, at step *i*, the MCMC is leaving the following unnormalized density invariant:

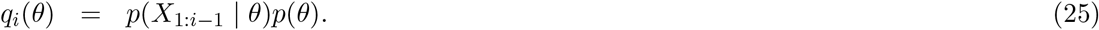

In the modified version, a family of reference distributions is introduced, *p_ϵ_*(*θ*), for 0 ≤ *ϵ* ≤ 1, such that, when *ϵ* = 0, *p*_0_(*θ*) is an estimate of the posterior distribution obtained based on a preliminary run, while, for *ϵ* = 1, *p*_1_(*θ*) reduces to the original prior *p*(*θ*). Then, at step *i*, the MCMC is targeting the following unnormalized density:

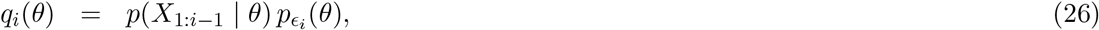

where *ϵ_i_* = min(1, 2(*i* − 1)/*n*). Thus, the family of reference distributions implements an interpolation between the empirical estimate of the posterior and the prior, starting under the empirical posterior distribution *p*_0_(*θ*) for *i* = 1, progressively moving toward the true prior for the first half of the dataset, reaching the prior when *i* > *n*/2 and then staying under the prior for the second half of the dataset. The reason why the interpolation is implemented only over the first half of the dataset is that the computation of the k-fold CV score requires to be under the original prior for all data points between *q* + 1 and *n*. Thus, with this design, all CV-scores such that the training set is at least half of the entire dataset can be computed based on this single run, by summing over the relevant segment, which is guaranteed to be contained within the second half. The details about the reference priors depend on the specific model and are given in the Supplementary Material (section 1.3).

#### Monte Carlo estimation of the wAIC

Practical estimation of the wAIC is relatively straighforward, in that the theoretical expectation and variance terms of equation 12 are simply replaced by their Monte Carlo counterparts (empirical mean and variance over the MCMC sample).

#### Bias estimation and correction

All Monte Carlo estimators presented above entail a step in which the expectation of a likelihood (or its inverse for CPO) is replaced by an arithmetic mean over the corresponding MCMC sample (equations 16 and 17 for nIS, 18 for CPO, 22 for sIS, and 12 for the wAIC). These Monte Carlo estimators are unbiased on the natural scale. However, the resulting scores are ultimately log-transformed. This transformation introduces a bias, which is negative for all estimators except for CPO, for which it is positive.

When the variance of the Monte Carlo estimator (on the natural scale) is small, the absolute value of the bias is, to good approximation, equal to half of the variance of the log-likelihood values over which the average is performed. In turn, the variance of the log-likelihood values can be estimated based on two independent MCMC runs. This approach was implemented systematically for all estimators, except for k-fold joint CV, for which the variance is too large for this approach to be usable in practice. The details are given in the Supplementary Material (section 1.4).

#### Implementation of nIS, sIS and CPO under the normal model

In the case of the normal model, it is possible to sample the parameter *θ* directly from the posterior distribution, *p*(*θ* | *X*_1:*i*_) for any *i* (see Supplementary Material, section 5). The Monte Carlo implemented for the normal model takes advantage of this property, by sampling *θ_t_*, *t* = 1..*T* (for nIS and CPO) or *θ_it_*, for *i* = 1..*n* and *t* = 1..*T* (for sIS) independently from the relevant posterior distribution.

#### Implementation of nIS, sIS and CPO under the phylogenetic models

In the case of phylogenetic models, the implementation of PhyloBayesMPI was taken as a starting point. The basic routines of MCMC sampling defining a cycle were left unchanged. Naive importance sampling was already implemented for joint k-fold CV, as a simple post-analysis routine that scans the MCMC chain (after burnin) and averages the likelihood scores over the run. This routine was augmented to also output the sitewise k-fold CV, according to the method described above. Similarly, LOO-CV and the wAIC are jointly computed based on another post-analysis routine. The sIS method requires more specific additions to the current implementation: essentially, defining and implementing the family of reference distributions that are necessary for the variance reduction approach described above and implementing the routines for adding sites during the MCMC.

### Data and general settings

For the phylogenetic analyses, two previously published empirical datasets were considered:

- EF2: a multiple sequence alignment of elongation factor 2 in 30 eukaryotic species (627 aligned positions), taken from Lartillot & Philippe (2006);
- Metazoa: a concatenation of genes (35371 aligned positions) across 35 metazoans, along with 2 choanoflagellates and 12 fungi for the outgroup (Philippe *et al*., 2005).

The specific details, such as data randomization and subsampling, or the detailed settings of the Monte Carlo computations, are given in the Supplementary Material (section 2).

## Results

### Comparing alternative measures of fit on a simple analytical example

The alternative measures of model fit that are considered in this work are marginal likelihoods, or equivalently Bayes factors, leave-one-out CV (LOO-CV) and k-fold CV (with *k* = 5 and based on independent randomizations of the dataset), the latter in two versions: joint and site-wise (see methods for details). Since they differ in their mathematical definition, these alternative measures of model fit have no reason to agree quantitatively, or even qualitatively, on specific real cases. To examine this point, and before getting into phylogenetic examples, the conceptual and numerical issues are illustrated using simulations under a simple multivariate normal model for which analytical results are available. In this subsection, only the conceptual issues (i.e. the differences in the exact mathematical measures of model fit) are considered, the numerical issues being examined in the next subsection.

The normal model considered here is a variant of the model originally due to Bartlett (1957). The simulated data consist of a series of *n* real vectors of dimension *p*, noted (*X_i_*)_*i*=1..*n*_, which are i.i.d. from a multivariate normal distribution of mean *θ*_*_ (also a p-vector) and of covariance matrix Σ = *σ*^2^*I_p_*, where *I_p_* is the identity matrix. The true mean *θ*_*_ used for simulation is chosen to be close to, but not equal to 0. Inference on these simulated data is conducted under two models.

In both models, the variance parameter *σ*^2^ is assumed known. Under model *M*_1_, the vector of means *θ* is fixed a priori to some value *θ*_0_, whereas it is re-estimated under model *M*_2_. Importantly, the aim is to represent a situation where model comparison is recruited for selecting the best approximating model, not the true model. Thus, what we want to formalize is a situation where the fixed parameter value *θ*_0_ defined by model *M*_1_ is never exactly true. Instead, *θ*_0_ may be viewed as a reasonably good proxy for the unknown true value *θ*_*_, and the question is just whether we can hope to get closer to *θ*_*_ by re-estimating *θ* on the dataset of interest, thus by using *M*_2_ rather than using *M*_1_. Accordingly, for *M*_1_, we set *θ*_0_ = 0 (which is thus different from, but close to, the true value *θ*\*). For model *M*_2_, we assume a normal prior on *θ*, of mean 0 and of covariance Σ_0_ = *δ*^2^*I_p_*. The hyper-parameter *δ* is chosen to be large, so as to implement a vague prior on *θ*. Of note, when *δ* → ∞, the prior becomes improper but the posterior reaches a well-defined limit. We wish to evaluate the relative fit of model *M*_2_ against *M*_1_ on a dataset of size *n*.

Data were more specifically simulated under the following settings: *p* = 300, *σ*^2^ = 10, *θ*_*_ = 0.1, and *n* varying from 100 to 10000. For model *M*_2_, two values were considered for the prior width, *δ*^2^ = 10 and *δ*^2^ = 1000. Then, the alternative measures of model fit were computed: marginal likelihood (Bayes factor), 5-fold cross-validation, both joint and site-wise, and leave-one-out cross validation. In all cases, the exact analytical values for the expected score of *M*_2_ relative to *M*_1_ were computed (Supplementary Material, section 5). The fit curves are displayed on Figure 1A, as a function of data size. Finally, an analytical formula is also available for the expected root mean squared error under the two models (using the posterior mean as the point estimate under model *M*_2_). This expected error, which is thus a frequentist risk, is displayed for the two models on Figure 1B as a function of data size.

**Figure 1:**
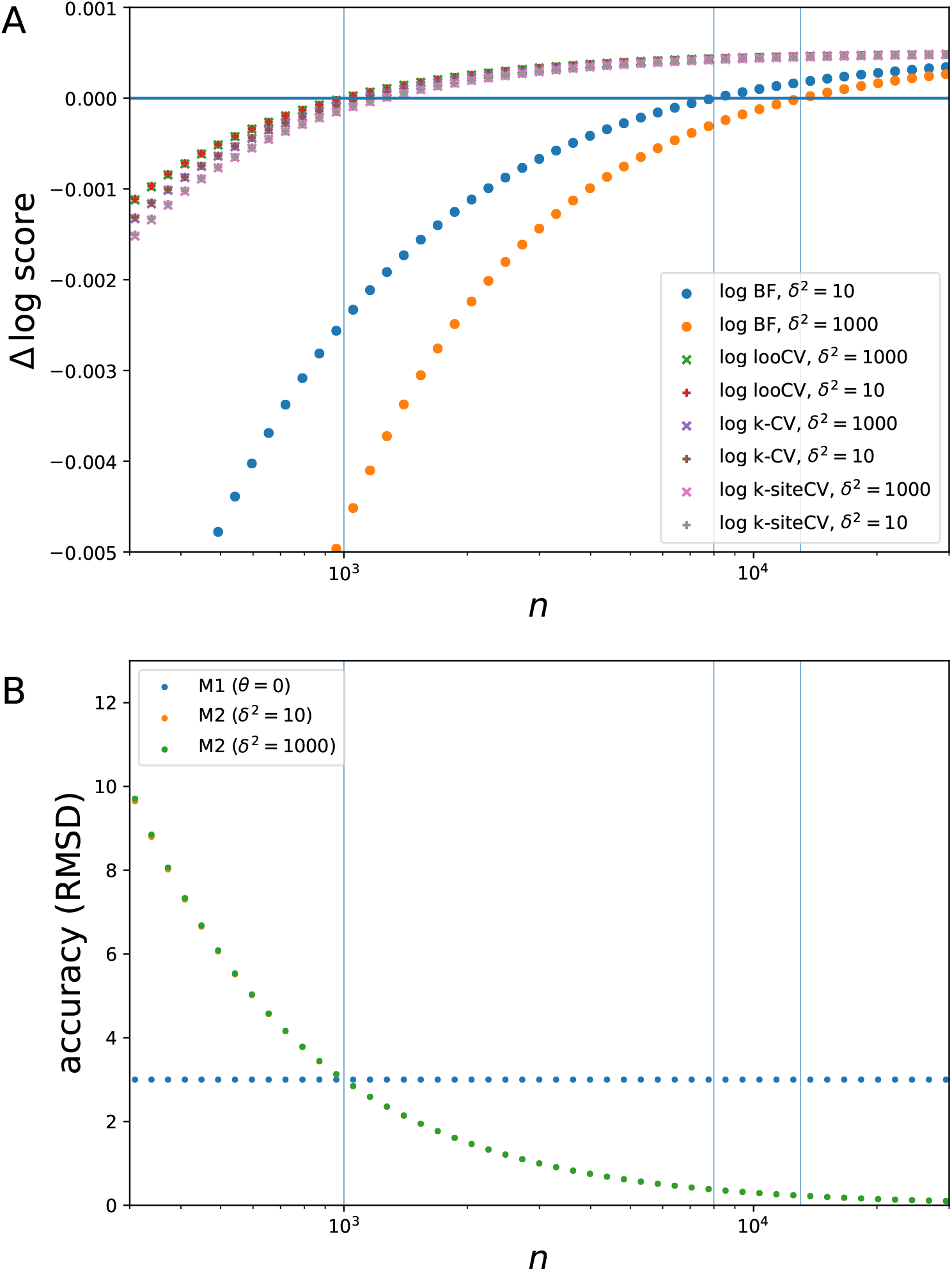
Theoretical fit of *M*_2_ relative to *M*_1_ (A) and mean squared estimation error (B) as a function of data size, under the normal model, and for two alternative priors (*δ*^2^ = 10 and 1000); vertical lines indicate the values of *n* for which the two models have same quadratic LOO-CV score or same marginal likelihood.

Several observations can be made from these experiments. First, for small data size, model *M*_1_ is more accurate than model *M*_2_. Model *M*_1_ is technically wrong (it assumes that *θ* = 0 whereas in fact *θ*_*_ > 0), however, for small data size, the estimation error under model *M*_2_ is much larger than the deviation between *θ*_*_ and 0, and thus it is indeed more reasonable to use *M*_1_ in that case. When *n >* 1000, on the other hand, *M*_2_ is more accurate than *M*_1_.

Second, by comparing the two panels of Figure 1, one can see that Bayes factors are clearly conservative. For instance, when *δ*^2^ = 10, it takes a dataset of at least 8000 observations for Bayes factors to show a preference for *M*_2_. Thus, between *n* = 1000 and *n* = 8000, Bayes factors are choosing a simple model that can be up to 5 times less accurate than the more complex alternative. This conservativeness is more pronounced under a broader prior (i.e. for larger *δ*). For *δ*^2^ = 1000, the cutoff at which Bayes factors switch to a preference for model *M*_2_ is slightly above *n* = 10000. Importantly, the posterior distribution is virtually the same for these two values of *δ*^2^, which shows that the differences in Bayes factors induced by the choice of the value of *δ* do not reflect any real-world difference, in terms of estimation. This conservative behavior of Bayes factors under vague priors is known as Jeffreys-Lindley’s paradox (Jeffreys, 1967; Lindley, 1957).

In contrast, the model chosen by CV appears to be more directly in proportion to estimation accuracy, with a cutoff very close to the tipping point (*n* = 1000) at which *M*_2_ starts to be more accurate than *M*_1_. In the details, k-fold CV appears a bit more conservative than LOO-CV, and site-wise k-fold CV is more conservative than both joint k-fold CV and LOO-CV. Although these differences are minor, they illustrate one potential problem with k-fold CV, namely, that it is not measuring the fit under the practically relevant data size. This limitation is inherent to cross-validation, but it is minimized in the case of leave-one-out, for which the training size is virtually indistinguishable from the practically relevant data size for even moderate values of *n*.

The asymptotic behavior of the alternative measures of fit explored here confirms these points. Up to an order 1/*n*, the logarithm of the Bayes factor (per site, i.e. 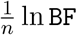) and the joint k-fold (Δ`cv`_*t*_), site-wise k-fold (Δ`cv`_*s*_) and leave-one-out (Δ`cv`_*l*_) cv scores of model *M*_2_ relative to *M*_1_ have the following expressions:

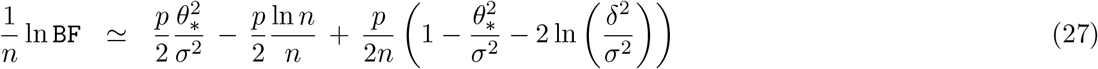

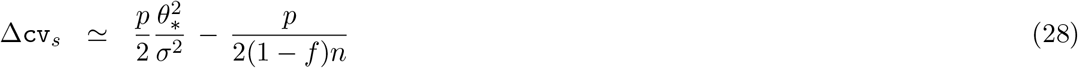

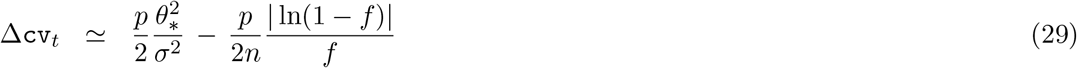

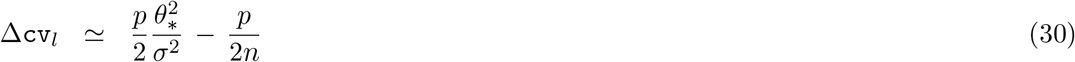

Of note, when the set-aside fraction *f* is small, then 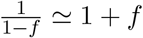, and 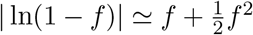, such that:

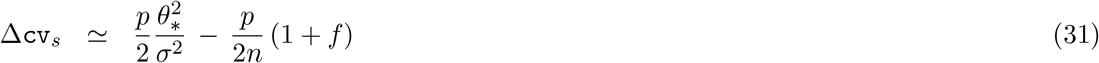

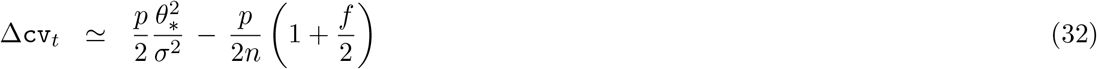

As for the asymptotic relative risk (i.e. difference in quadratic error between model *M*_1_ and *M*_2_, normalized here by 2*σ*^2^), it is, up to terms in 1/*n*:

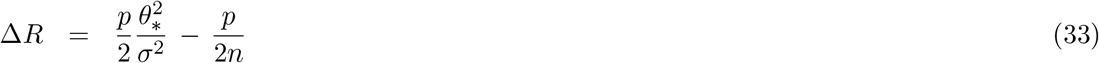

All these equations can be obtained as simple first-order developments of the exact analytical expressions that are provided in section 5 of the Supplementary Material.

From these equations, several observations can be made. On one hand, for sufficiently large *n*, all terms except the first vanish. Since this first term is identical, all measures will eventually agree and will all choose *M*_2_ (as also shown by equation 15). This asymptotic agreement is visible on Figure 1A. Also visible on the figure is the slower convergence, in ln *n/n*, for the log Bayes factor, whereas it is in 1/*n* for cross-validation under all settings. Of note, this mirrors the penalties of the BIC and the AIC, respectively.

Conversely, however, for fixed *n*, and considering increasingly vague priors by letting *δ* go to infinity, the log Bayes factor is ill-behaved, since its last term goes to −∞. In other words, for a given data size, and for arbitrary large *θ*_*_, then, provided that the prior is sufficiently broad, BF will nevertheless prefer *M*_1_ – and this, in spite of the arbitrary large risk that this might entail. In contrast, CV measures are all well-behaved and are insensitive to *δ*. When the set-aside fraction *f* is small, the two versions of k-fold CV are slightly more conservative than LOO-CV (that is, they have a slightly stronger penalty), the joint version being intermediate between LOO-CV and the site-wise version, as seen on Figure 1A. Finally, LOO-CV is asymptotically equal to the difference in quadratic estimation error between the two models. In other words, asymptotically, LOO-CV is exactly selecting the model that gives the most accurate estimate. However, this last point is not a general result. Instead, it is a consequence of the fact that a spherical covariance matrix was used in the model. For general covariance structures, the LOO-CV score is asymptotically equal to another relative quadratic risk, computed under the metric defined by the covariance matrix, that is, based on the quadratic error defined as *L*_Σ_ = ^*t*^(*θ* − *θ*_*_)Σ^−1^(*θ* − *θ*_*_). This metric essentially gives less weight to the errors made on those components of *θ* for which the likelihood is less informative.

A final point not quantitatively explored here but worth noting: when *θ*_*_ = 0, that is, when *M*_1_ is the true model, all CV methods considered here are asymptotically inconsistent, in the sense that the probability of choosing *M*_1_ does not converge to 1 for large *n* (Shao, 1993). However, whenever CV chooses *M*_2_, it will then estimate a value for *θ* very close to its true value 0 (up to a quadratic error in 1/*n*), such that the error in model selection will have a negligible impact on estimation accuracy. In other words, CV is not formally consistent, but it is effectively consistent, in the sense that the selected model is asymptotically equivalent to the true model in the Kullback-Leibler metric. Conversely, any method trying to be asymptotically formally consistent will have to be more strongly penalizing than LOO-CV. However, since LOO-CV is asymptotically optimal in estimation accuracy when the true model is not *M*_1_, such a method will be suboptimal for selecting the best approximating model in that case. The two goals of model selection, best approximation or true model identification, are thus mutually incompatible (Shibata, 1986).

### Numerical approaches: accuracy and computational considerations

In this section, the question of the numerical evaluation of the alternative measures of model fit considered above is explored, again in the normal case, for which the numerical estimates can be directly assessed against the analytically available value. The Monte Carlo approaches that are used here are all variations on importance sampling: naive importance sampling (nIS) for both joint and site-wise k-fold CV, sequential importance sampling (sIS) for joint k-fold CV and marginal likelihoods, and the cross-predictive ordinate (CPO) approach for LOO-CV (see methods for details). The results are presented in Table 1.

**Table 1:** Numerical estimates of the fit of *M*_2_ (relative to *M*_1_) under various criteria and numerical approaches for the normal model example. See text for details.

| method | dim ($p$) | sample size | | model fit ($\Delta$ log score) | | | bias | | error | |
| --- | --- | --- | --- | --- | --- | --- | --- | --- | --- | --- |
|  |  | nominal | ESS | true | est.<sup>1</sup> | deb.<sup>2</sup> | true<sup>3</sup> | est.<sup>4</sup> | raw<sup>5</sup> | deb.<sup>6</sup> |
| k-CV (nIS) | 100 | $10^4$ | 6.49 | 0.02 | 0.017 | 0.018 | -0.006 | -0.001 | 0.009 | 0.009 |
| | 100 | $10^6$ | 24.20 | 0.02 | 0.021 | 0.021 | -0.002 | -0.000 | 0.004 | 0.004 |
| | 300 | $10^4$ | 2.26 | 0.08 | 0.010 | 0.011 | -0.065 | -0.001 | 0.067 | 0.066 |
| | 300 | $10^6$ | 2.95 | 0.08 | 0.033 | 0.035 | -0.042 | -0.001 | 0.043 | 0.042 |
| | 1000 | $10^4$ | 1.52 | 0.24 | -0.159 | -0.157 | -0.394 | -0.002 | 0.395 | 0.393 |
| | 1000 | $10^6$ | 1.74 | 0.24 | -0.091 | -0.090 | -0.327 | -0.002 | 0.328 | 0.326 |
| k-site-CV (nIS) | 100 | 10 | 9.01 | 0.02 | 0.008 | 0.015 | -0.007 | -0.006 | 0.012 | 0.010 |
| | 100 | $10^3$ | 883.02 | 0.02 | 0.015 | 0.015 | -0.000 | -0.000 | 0.001 | 0.001 |
|  | 300 | 10 | 7.55 | 0.06 | 0.035 | 0.055 | -0.020 | -0.020 | 0.026 | 0.017 |
| | 300 | $10^3$ | 689.11 | 0.06 | 0.055 | 0.056 | -0.000 | -0.000 | 0.002 | 0.002 |
|  | 1000 | 10 | 5.15 | 0.16 | 0.063 | 0.127 | -0.101 | -0.064 | 0.106 | 0.050 |
| | 1000 | $10^3$ | 302.75 | 0.16 | 0.162 | 0.163 | -0.002 | -0.001 | 0.004 | 0.004 |
| k-CV (sIS) | 100 | 10 | 9.09 | 0.02 | 0.018 | 0.023 | -0.006 | -0.006 | 0.008 | 0.006 |
| | 100 | $10^3$ | 894.89 | 0.02 | 0.023 | 0.023 | -0.000 | -0.000 | 0.001 | 0.001 |
|  | 300 | 10 | 7.74 | 0.08 | 0.057 | 0.077 | -0.018 | -0.020 | 0.020 | 0.010 |
| | 300 | $10^3$ | 716.99 | 0.08 | 0.075 | 0.075 | -0.000 | -0.000 | 0.001 | 0.001 |
|  | 1000 | 10 | 5.39 | 0.24 | 0.153 | 0.236 | -0.082 | -0.083 | 0.084 | 0.023 |
| | 1000 | $10^3$ | 341.29 | 0.24 | 0.234 | 0.235 | -0.001 | -0.001 | 0.003 | 0.002 |
| BF (sIS) | 100 | 10 | 8.91 | -0.33 | -0.339 | -0.331 | -0.008 | -0.007 | 0.008 | 0.003 |
| | 100 | $10^3$ | 871.21 | -0.33 | -0.331 | -0.331 | -0.000 | -0.000 | 0.000 | 0.000 |
|  | 300 | 10 | 7.39 | -1.00 | -1.020 | -0.995 | -0.024 | -0.025 | 0.024 | 0.005 |
| | 300 | $10^3$ | 663.74 | -1.00 | -0.996 | -0.996 | -0.000 | -0.000 | 0.001 | 0.001 |
|  | 1000 | 10 | 4.98 | -3.30 | -3.420 | -3.310 | -0.116 | -0.110 | 0.117 | 0.013 |
| | 1000 | $10^3$ | 277.90 | -3.30 | -3.306 | -3.304 | -0.002 | -0.002 | 0.002 | 0.001 |
| LOO-CV (CPO) | 100 | 10 | 9.18 | 0.03 | 0.035 | 0.030 | 0.005 | 0.005 | 0.006 | 0.003 |
| | 100 | $10^3$ | 904.89 | 0.03 | 0.030 | 0.030 | 0.000 | 0.000 | 0.000 | 0.000 |
|  | 300 | 10 | 7.93 | 0.09 | 0.103 | 0.086 | 0.018 | 0.017 | 0.019 | 0.006 |
| | 300 | $10^3$ | 741.22 | 0.09 | 0.085 | 0.085 | 0.000 | 0.000 | 0.001 | 0.001 |
|  | 1000 | 10 | 5.60 | 0.30 | 0.374 | 0.302 | 0.073 | 0.072 | 0.074 | 0.012 |
| | 1000 | $10^3$ | 376.91 | 0.30 | 0.302 | 0.301 | 0.001 | 0.001 | 0.001 | 0.001 |

<sup>1</sup>: estimated; <sup>2</sup>: debiased; <sup>3</sup>: true bias (difference between estimated and true fit); <sup>4</sup>: estimated bias (based on the Monte Carlo variance of log-likelihoods, see methods) <sup>5</sup>: true error of raw estimator; <sup>6</sup>: true error of debiased estimator

#### Naive importance sampling for k-fold CV

Naive importance sampling simply consists of averaging the likelihood of the data points of the validation set (either jointly or separately, for joint or site-wise k-fold CV respectively) over a sample of parameter configurations drawn from the posterior distribution under the training set. When applied to joint k-fold CV, nIS works well for low dimension (*p* = 10, 30 or 100) but its performance progressively degrades as the dimension of the model increases (Table 1). For large dimension *p >* 300, a substantial downward bias is observed. In the case of *p* = 1000, the bias is sufficiently strong to change the qualitative outcome of model selection, leading to an apparent CV score in favor of model *M*_1_, whereas model *M*_2_ has mathematically a higher CV score. As expected, increasing the size of the Monte Carlo sample can improve the situation, although very moderately. Under the highest dimensions considered here, it seems that it would take samples of very large size, well above 10^6^, in order to reduce the bias down to reasonable values.

A key statistic that is able to issue a warning about the reliability of the estimation in the present case is the effective sample size (ESS). The ESS is a function of the variance of the importance weights (formally defined in section 1 of the Supplementary Material), such that the ESS is close to 1 when a single point of the sample has an overwhelming contribution to the Monte Carlo average (essentially, the point of the sample that happens to have the highest likelihood). In the case of k-fold CV, for high dimensions, the ESS is indeed close to 1, indicating that the estimator is fundamentally unreliable.

In contrast to what is observed for joint k-fold CV, nIS works well on site-wise k-fold CV (Table 1). This is due to the fact that the single-observation likelihood is much less peaked than the joint likelihood of multiple observations. As a result, the variance of the log-likelihood score under the posterior distribution is small. Of note, for small MCMC sample size (10 samples per site), the total bias of the estimator in log scale can be non-negligible. On the other hand, because this bias is a sum of many small contributions (one for each data point), each of which has a large ESS and therefore a small variance, it can be estimated based on a linear approximation relating it to the variance observed across two independent runs (see methods). As a result, it can be worth de-biasing the estimator; although not perfect, doing this does increase the overall accuracy, quite substantially (Table 1).

#### A sequential importance sampling approach for k-fold CV and BF

As a way to overcome the limitations of naive IS, an alternative approach was implemented, based on sequential importance sampling (sIS, see methods). When applied to the multivariate normal problem, sIS gives a more reliable estimate of the joint k-fold CV score over the whole range of model dimensionalities considered here (Table 1). The estimate of the log marginal likelihood returned by sIS is also reliable, for both small and large dimensions. Here again, as in the site-wise case, the total bias of the estimators can be substantial for small Monte Carlo sample size, but is itself well-estimated. This sIS approach, however, is expensive – even more expensive for CV than for marginal likelihood, since CV requires running sIS ideally over a large number of randomized replicates of the original dataset, whereas only two runs on the original non-randomized alignment are needed for the marginal likelihood.

#### Leave-one-out cross validation using cross-predictive ordinates

An estimate of the LOO-CV score can be obtained very efficiently, based on a standard MCMC run under the posterior distribution, using the CPO approach (Gelfand *et al*., 1992; Chen *et al*., 2012; Lewis *et al*., 2014). The CPO method gives accurate estimates of the LOO-CV score (Table 1). Here again, the bias can be substantial for small sample size but is well estimated.

Altogether, naive IS works well for site-wise k-fold CV, but does not work well for joint k-fold CV. Both joint k-fold CV and Bayes factors require computationally intensive MCMC approaches, such as sIS. Finally, the CPO approach represents a reliable and computationally efficient method for estimating the LOO-CV score.

### An empirical example using a single-gene alignment

The various scores and numerical methods for computing them were then implemented in PhyloBayes (Lartillot *et al*., 2009, 2013). In a phylogenetic context, it is natural to use the individual columns of the multiple sequence alignment as the individual data points. For the rest, the implementation of all of the methods is relatively straightforward, based on the already existing MCMC routines. All of these estimators were then jointly examined, in the context of a global comparison between alternative site-homogeneous and site-heterogeneous models on an empirical alignment (elongation factor 2 in 30 eukaryotic species, 627 aligned positions, see Methods). The models under comparisons are the Poisson model (exchangeabilities between amino-acids all equal to 1), the empirical matrices WAG (Whelan & Goldman, 2001) and LG (Le & Gascuel, 2008), and finally, the CAT-Poisson model (Lartillot & Philippe, 2004), in two alternative versions that differ in the base distribution used for the Dirichlet process over the amino-acid frequency vectors: either a uniform (fix-hyper) or a general (free-hyper) Dirichlet distribution whose hyperparameters are then also estimated. The latter is the version of the CAT model proposed by default by PhyloBayes. The results are presented in Table 2.

**Table 2:** Numerical estimates of the fit of alternative substitution models for the elongation factor alignment. All measures of fit are on a logarithmic scale and on a per-site basis. Entries in bold correspond to top-ranking models

| model | 5-fold CV |  |  |  |  |  | BF |  | LOO-CV |  |
| --- | --- | --- | --- | --- | --- | --- | --- | --- | --- | --- |
| | nIS ($10^3$) | | nIS ($10^4$) | | sIS | | sIS | | CPO | |
|  | fit | (ESS) | fit | (ESS) | fit | (ESS) | fit | (ESS) | fit | (ESS) |
| Poisson | -30.89 | ( 3.3) | -30.88 | ( 6.7) | -30.87 | (28.4) | -31.49 | (28.3) | -31.35 | (856.5) |
| WAG | -28.18 | ( 5.7) | -28.17 | ( 8.3) | -28.16 | (28.3) | -28.82 | (28.2) | -28.68 | (864.2) |
| LG | -27.96 | ( 5.1) | -27.95 | ( 6.2) | -27.94 | (28.3) | -28.61 | (28.3) | -28.47 | (864.5) |
| GTR | -27.92 | ( 1.8) | -27.89 | ( 2.3) | -27.79 | (31.6) | -28.65 | (31.8) | -28.31 | (675.2) |
| CAT-fix-hyper | **-27.54** | ( 1.8) | **-27.51** | ( 2.5) | -27.34 | (33.6) | -28.10 | (36.8) | -27.80 | (658.1) |
| CAT-free-hyper | -27.66 | ( 1.3) | -27.59 | ( 1.5) | **-27.18** | (37.1) | **-28.04** | (36.0) | **-27.67** | (552.0) |

First, concerning k-fold CV, nIS and sIS (which are two alternative estimators of the same mathematical quantity) agree with each other on simple models such as Poisson or WAG, but not for more complex models such as CAT-Poisson. The ESS clearly suggests that, here also, as in the normal case, nIS is being unreliable. In one case, this leads to a different qualitative answer as to which model is best fitting. Thus, nIS gives an apparently higher joint k-fold CV score for the version of the CAT model that uses a uniform base distribution (fix-hyper), whereas sIS says that the version with a general Dirichlet distribution (free-hyper) has a higher fit. Of note, fix-hyper is a constrained version of the free-hyper model, assuming a uniform base distribution. The posterior estimate of the base distribution under the unconstrained model, however, is very far from uniform: the posterior mean estimate and 95% credible interval for the mean concentration parameter of the Dirichlet distribution is *β* = 5.8 (4.8, 6.8), whereas it is equal to 20 for the uniform distribution over the simplex. This gives an additional argument, independent from the ESS, suggesting that nIS is giving a wrong answer in the present case.

Second, concerning the alternative measures of model fit: k-fold CV and LOO-CV qualitatively agree with each other. The differences in predictive fit between models of widely differing dimension is slightly more pronounced for LOO-CV than for k-fold CV. Thus, for instance, the fit of CAT-Poisson (with free hyper-parameters) relative to LG is estimated at 0.80 according to LOO-CV, versus 0.76 according to k-fold CV. This mirrors the pattern seen previously in the normal case (Figure 1), namely, that k-fold CV, because it is trained on a subset of the data, tends to underestimate the predictive fit of more complex models, relative to simpler ones. This effect is small, however, much smaller than the numerical error of nIS on joint k-fold CV (which gives a relative fit of 0.36, instead of 0.76, for CAT-Poisson versus LG).

If LOO-CV and k-fold CV qualitatively agree with each other, on the other hand, they differ somewhat from the Bayes factor, which tends to be more conservative. In one case, BF and CV give a qualitatively different outcome, concerning the choice between GTR and empirical matrices: whereas all CV methods choose GTR, BF gives a higher score to the LG model on this EF2 dataset.

### LOO-CV, BF and estimation accuracy in a phylogenetic context

The experiment above on EF2 suggests that BF can sometimes disagree with cross-validation on real cases. To further investigate this point, another experiment was conducted, consisting of comparing LG and GTR on increasingly large subsets of an empirical supermatrix of 35 metazoan species (Philippe *et al*., 2005), using either BF or LOO-CV. The k-fold CV approach was not considered, owing to its computational cost. Of note, here as above (Table 2), the prior on the renormalized exchangeabilities (constrained to sum to 1) of the GTR model is uniform, thus uninformative.

The results of this experiment are summarized in Figure 2. For sufficiently large datasets, BF and LOO-CV both favor the GTR model over LG, while for smaller datasets, the LG model tends to be favored. This point is expected, and confirms that, for sufficiently large data size, there is an opportunity for getting better estimates of the relative exchangeabilities than those proposed by LG. However, if both methods agree on this dichotomy between small versus large datasets, they differ substantially concerning the exact cutoff, in terms of data size, at which they switch from LG to GTR: whereas BF favors GTR over LG only starting from alignments made of more than 600 sites, LOO-CV does so for datasets as small as 200 sites.

**Figure 2:**
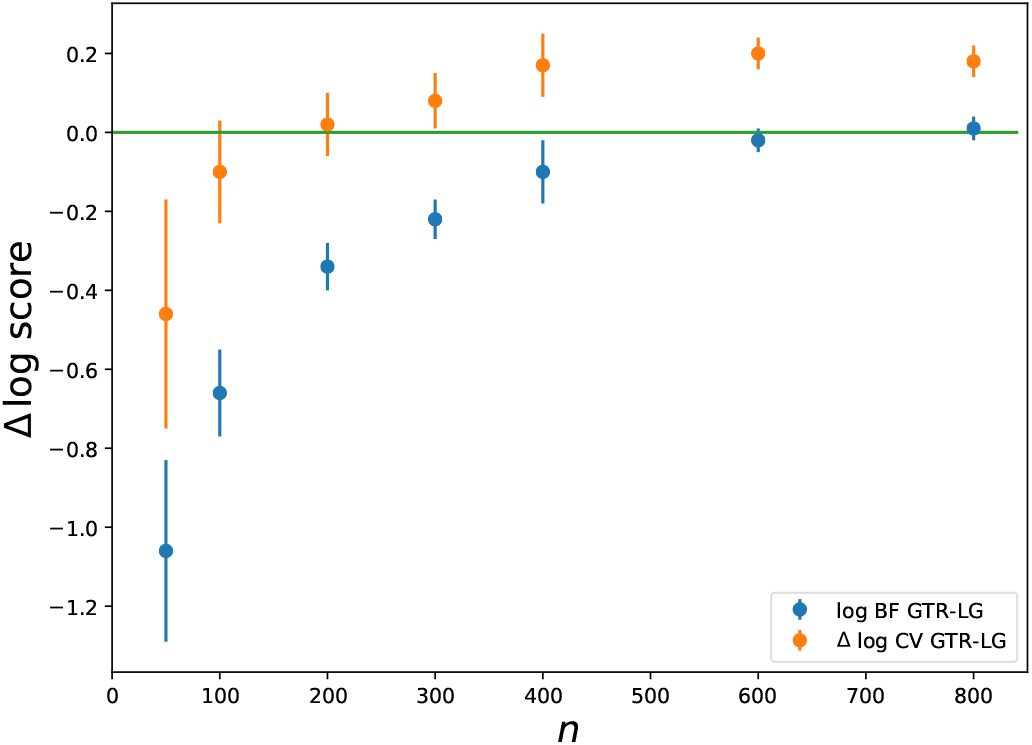
Fit of GTR, relative to LG, as a function of data size (number of aligned positions), on empirical data (10 random jackknife subsamples of the metazoan dataset), using BF and LOO-CV. Error bars: standard deviation across jackknife replicates.

The analytical results presented above under the normal model suggested that cross-validation is more in phase with estimation accuracy than Bayes factors. To investigate whether this conclusion is also valid in the present case, the following simulation experiment was conducted. First, data were simulated under LG and under empirically calibrated branch lengths and parameter values, using the posterior predictive formalism and with the metazoan alignment as a template (see Supplementary Material, section 2). Then, model selection was implemented, between JTT and GTR. Importantly, the true model (LG) was not included in the set of models being compared. This omission is meant to represent the fact that, in reality, the true exchange rates (or, more accurately, the asymptotic exchange rates, i.e. the ones that would be eventually estimated on a sufficiently large alignment obtained from this empirical source) are not equal to any of the empirical models that are available. The difference between JTT and LG is thus meant as a representation, in our simulation experiment, of the difference between LG and the true exchange rates in the empirical experiment.

The Bayes factors and cross-validation scores obtained on these simulated data (Figure 3A) reproduce the pattern observed on the empirical data (Figure 2) as a function of data size, with GTR being ultimately favored by both BF and LOO-CV, although for a larger cutoff data size for BF (700) than for LOO-CV (200). Of note, both the cutoffs and the absolute fit values are very similar to those obtained on the original empirical experiment (Figure 2), suggesting that the simulation experiment is mimicking the true empirical situation relatively well.

**Figure 3:**
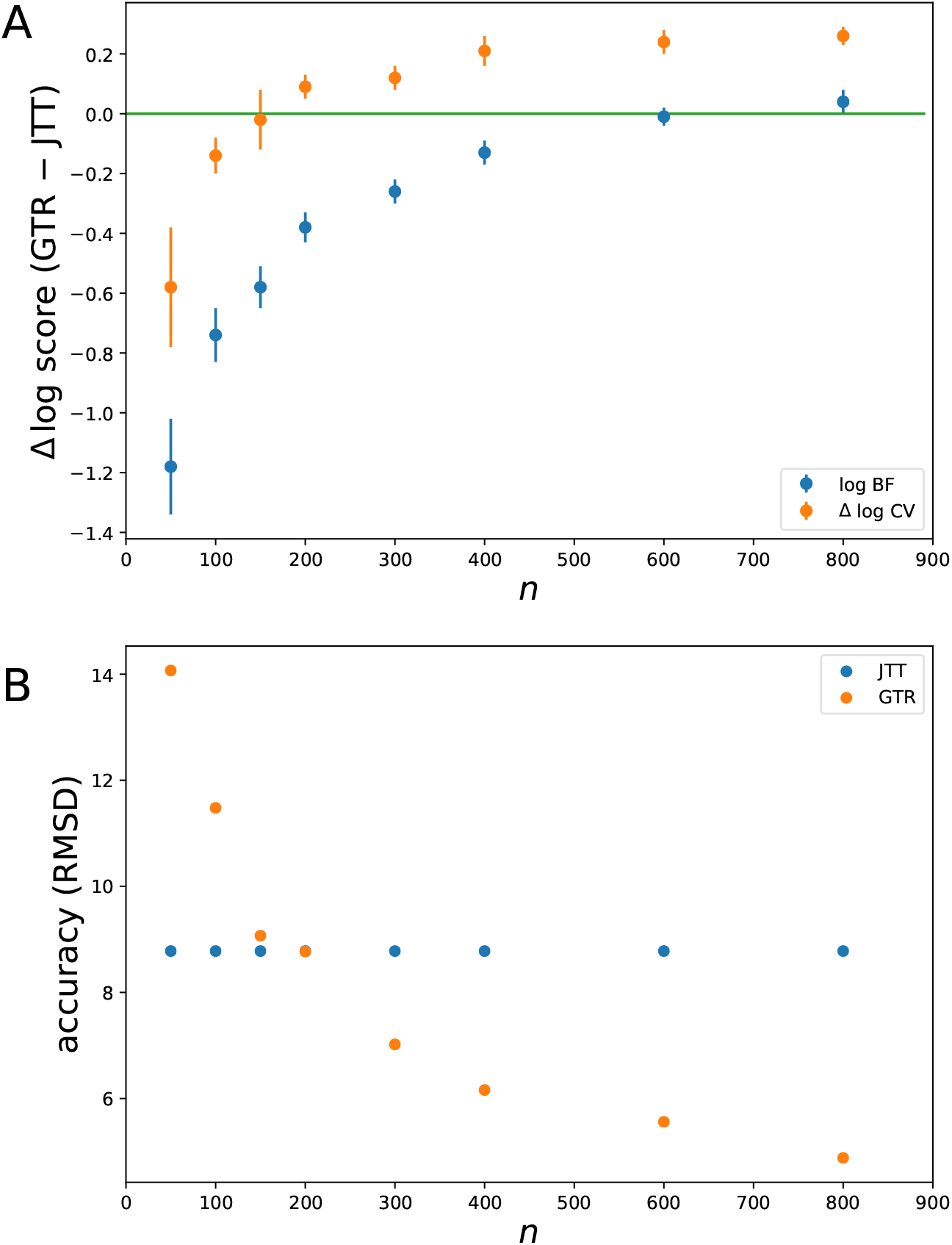
Fit of GTR, relative to JTT, as a function of data size under BF and LOO-CV (A), and mean quadratic error on relative exchangeability estimates (B) on data simulated under the LG model (using the metazoan dataset as a template). Error bars: standard deviation across 4 simulation replicates.

Along with model fit, the error (RMSD) in the estimation of the relative exchange rates was also quantified (Figure 3B). In the case of the JTT model, this error is trivially constant (quadratic deviation between JTT and LG). For the GTR model, this error decreases with data size. On sufficiently small alignments, on the other hand (smaller than 200), the estimation error under GTR can be larger than the difference between JTT and the true exchange rates (LG). Thus, for small alignments, we are in a case where the most accurate model is in fact JTT, and this, in spite of the fact that JTT is not the true model.

Finally, comparing RMSD with both BF and LOO-CV shows that LOO-CV provides a good predictor of which model is more accurate for parameter estimation, with a cutoff at around 200 aligned positions. BF in contrast, imposes a stronger penalty and still chooses JTT for datasets up to 600 sites, thus well within the regime of alignment size where GTR is in fact already returning a substantially more accurate estimation. Transposing these observations to the empirical case, this suggests that LG is in fact not so good and GTR is better, even for small alignments of about 200 sites and 50 taxa. It also confirms the point already demonstrated on the normal case, namely that LOO-CV gives a more reliable predictor of estimation accuracy than BF.

### Asymptotics of LOO-CV and the widely applicable information criterion (wAIC)

The asymptotic equivalence between the wAIC and LOO-CV (Watanabe, 2010*a*) was empirically assessed by conducting a scaling experiment, consisting of randomly subsampling a large phylogenomic dataset and plotting the fit (LOO-CV and wAIC) of the GTR, CAT-Poisson and CAT-GTR models (relative to LG), as a function of data size (Figure 4).

**Figure 4:**
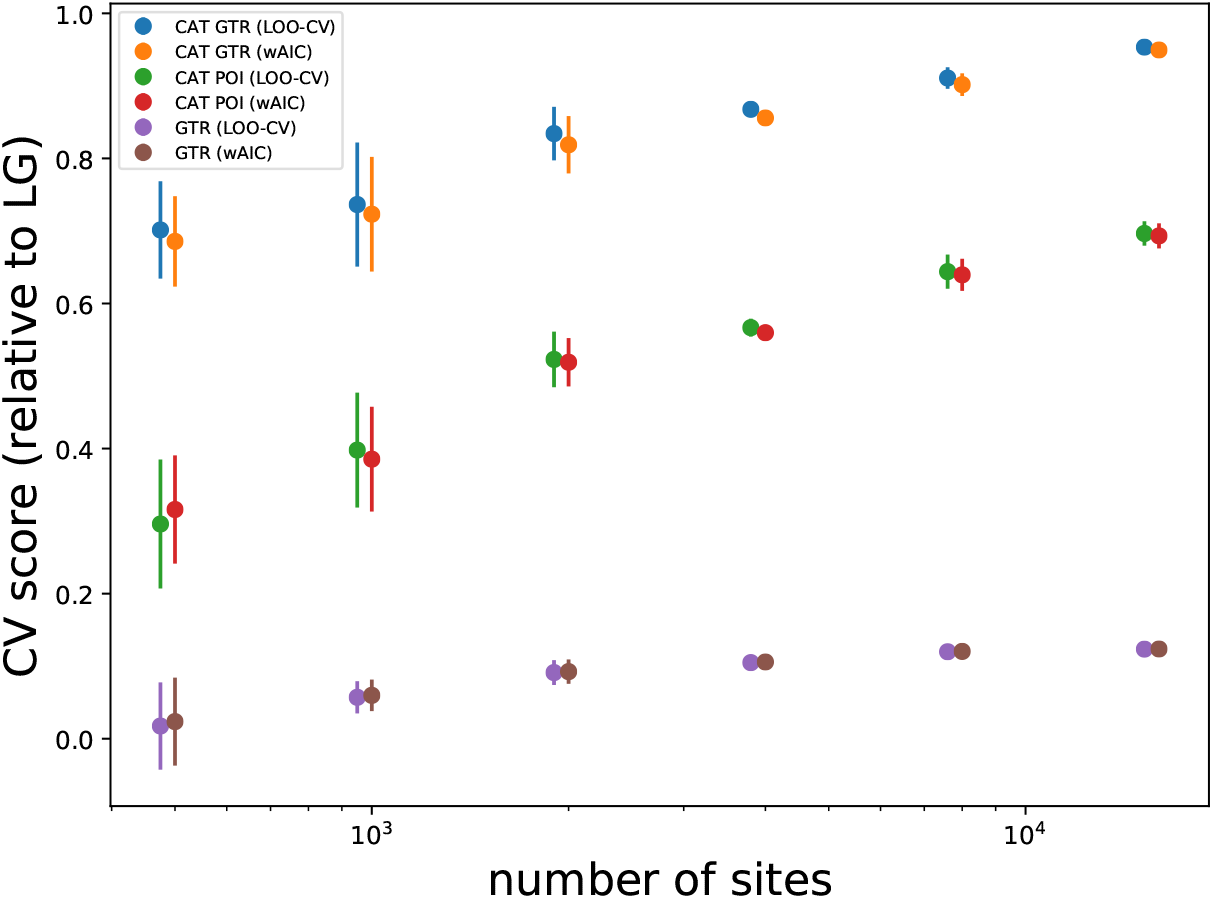
LOO-CV and wAIC estimates (de-biased) for the GTR, CAT-Poisson and CAT-GTR models (relative to LG), as a function of data size (number of aligned positions), on empirical data (metazoan dataset). Error bars: standard deviation across 4 jackknife replicates.

Overall, LOO-CV and the wAIC give very similar results. The discrepancy between them decreases as the data size becomes larger, giving nearly indistinguishable numerical estimates for the largest data sizes considered here. Even for smaller data size, the difference between LOO-CV and the wAIC is visible but small compared to the difference in fit between the models.

The numerical stability and the accuracy of the two estimators were investigated in more detail, using a combination of several approaches. First, several statistics were monitored: the estimated bias and standard deviation, and the distribution of effective sample sizes (ESS) across sites. Second, the impact of MCMC sample size was assessed, by using either the full sample (of size *T* = 1000) or by thinning down to *T* = 100 samples from the posterior. On practical grounds, thinning represents a particularly attractive option in the case of mixture models, for which the numerical evaluation of the likelihood, as a sum over mixture components, is computationally intensive and turns out to be the limiting factor for estimating the fit. Third, as an alternative to CPO, the Pareto smoothed importance sampling estimator PS-LOO-CV (see methods and Vehtari *et al*., 2016) was also used, and the associated quality checks were monitored.

Both estimators, of LOO-CV and of wAIC, tend to be numerically more stable for larger MCMC sample size, but also for larger data size (Supplementary Table 1). For smaller data size (*n* ≤ 1000), there are warnings about a fraction of sites with critically low ESS (or with high values of 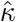 according to PS-LOO-CV), in particular for the more complex model (CAT-Poisson). Compared to LOO-CV, the wAIC tends to show better quality checks based on ESS and is less sensitive to variation in Monte Carlo sample size.

The estimators of LOO-CV, PS-LOO-CV and the wAIC were further benchmarked against an independent approach. The scaling experiment conducted here, through it reliance on a large dataset representing a proxy for the population, offers an opportunity for a more direct operational implementation of the ideal target of cross-validation and the wAIC, namely, by training the model on an independent data set of same size, before validating it (using the sitewise cross-validation approach) on the dataset of interest. This approach, hereafter called 2-step CV, is numerically more stable and shows an opposite numerical bias, compared to LOO-CV. For that reason, it represents a useful point of comparison.

This more extensive benchmark confirms the observations made above. In the case of GTR, all estimators, PS-LOO-CV, LOO-CV, wAIC and 2-step CV, are virtually indistinguishable over the whole range (Supplementary Figure 1). In the case of CAT-Poisson, larger estimation errors are observed for small data size (*n* ≤ 1000). For large data size, on the other hand, all estimators return very similar estimates. PS-LOO-CV does not seem to bring an improvement over LOO-CV and shows a stronger positive bias. Finally, de-biasing the estimators is helpful, in particular for small MCMC sample size, such that LOO-CV, the wAIC and 2-step-CV end up being closer to each other after bias correction, showing differences that are small compared to the difference between models and to the sampling variance, and this, even based on thinned sample of 100 points (see Supplementary Material, section 3, for a more extended analysis).

## Discussion

In many respects, model comparison and model selection in Bayesian inference is still an open problem. Conceptually, in spite of a large literature on the question, a general agreement on the guiding principles has not yet been achieved. Computationally, numerical inaccuracies are surfacing regularly. The present work attempts to bring a few points of clarification, along with a correction concerning the numerical accuracy of a previously introduced importance sampling k-fold CV approach. In the end, some recommendations are suggested for improving both reliability in model selection and computational accuracy and efficiency.

The main conclusions are as follows. As suggested previously (Gelfand *et al*., 1992; Bernardo & Smith, 1994; Konishi & Kitagawa, 2007), Bayes factors are inadequate for selecting the best-approximating model, and cross-validation appears to be more adequate for this purpose. Among CV methods, LOO-CV stands out as the best choice, both statistically and computationally. It also has a clear asymptotic connection with information criteria, and more specifically with the widely applicable (or Watanabe-Akaike) information criterion (wAIC Watanabe, 2009). For large datasets, wAIC is easily implemented and offers a good complement to LOO-CV.

### Problems with marginal likelihoods under vague priors

One first fundamental reason for the conservative behavior of marginal likelihoods in the present case is the use of a vague prior over the model-specific parameters. From the experiments presented here, and more generally on conceptual grounds, marginal likelihoods do not represent a meaningful measure of model fit under a prior that is meant to be uninformative. This is particularly apparent in the case of the normal model. Under this model, when the prior over the unknown mean *θ* is uniform over the entire real line, and thus improper, the posterior distribution is well-defined, but the marginal likelihood is infinite. This problem has been known for a long time (Gelfand *et al*., 1992; Jeffreys, 1967; Lindley, 1957), and it has already been noted in the context of phylogenetic inference that marginal likelihoods and Bayes factors should not be used with improper priors (Baele *et al*., 2012*b*). However, making the prior technically proper but still effectively uninformative does not solve the problem. This is again clear in the case of the normal model, for which model selection based on the marginal likelihood can be made arbitrarily stringent against the more complex model by playing on its width parameter *δ* – and this, in spite of the fact that the posterior distribution is virtually unaffected (Figure 1b, compare red and green dots). This problem is also well illustrated by the comparison between JTT and GTR. In that case, the prior over the relative exchangeabilities of the GTR model is proper, but non informative, and the marginal likelihood is unduly biased in favor of JTT.

A reasonable operational consistency requirement in the context of best-approximating model selection would be that the criterion used for selecting models should give essentially identical scores to models that give essentially identical posterior distributions. Obviously, marginal likelihoods do not fulfill this consistency requirement. They are are notoriously sensitive to the prior – and more so than the posterior distribution itself. In contrast, cross-validation, and its asymptotic equivalent given by information criteria such as the wAIC, are by construction dependent on the prior only through the posterior distribution. Thus, they are guaranteed to be operationally consistent.

Importantly, all this does not imply that using uninformative priors is in itself problematic. Uninformative priors do have a good theoretical justification, as a bet-hedging strategy, whose aim is to minimize the worst case error over all possible values the unknown parameter might have (Berger, 1985). As such, they are generally proposed as default priors, meant to guarantee some robustness in the context of automatic application of the inference method to an arbitrary series of practical cases (Berger, 2006). They are thus particularly useful as routine priors, in particular for the global parameters of the model, which are at the top of the hierarchy. However, model selection methods should then be compatible with these priors.

### Cost of learning versus accuracy, and the two aims of model selection

Intuitively, another fundamental problem of marginal likelihoods in the present context is that they penalize models in proportion to how much information information has been extracted from the data and how well it agrees with the prior (essentially, the relative width and position of the posterior, compared to the prior), and not in proportion to how accurately this information has been learned. Yet, for selecting the best-approximating model, only the second point is relevant. The penalty induced by cross-validation, on the other hand, is directly and exclusively related to how well the fitted model predicts new data. As a result, it is more directly related to the accuracy of the end result of the estimation, leaving out any consideration about the total cost of parameter fitting.

The sequential importance sampling formalism gives another intuition of the same idea. With sIS, the logarithm of the marginal likelihood is obtained by starting from the prior, adding sites incrementally and summing up their individual contributions. By the chain rule (equation 21), the overall marginal likelihood score is a sum over the total learning curve, and as a result, it penalizes models in proportion to the total learning work done upon going from the prior to the posterior. In contrast, leave-one-out cross validation considers only the last step of the procedure and therefore penalizes in proportion to the marginal surprise of the last data point (taken as a proxy for the average future observation).

On the other hand, the total cost of fitting and, more generally, the sensitivity of the marginal likelihood to the prior, is potentially relevant for hypothesis testing. For instance, an alternative hypothesis may explain the data better than does the null, but only under an effect size that is very small compared to the typical effect sizes that would be a priori expected if the alternative were true. This a priori unlikely event will represent a cost that marginal likelihoods will incorporate in their evaluation of the fit, making them more inclined to select the null hypothesis in that case. Marginal likelihoods will thus be useful in a hypothesis testing context, although this requires careful design of the priors over effect sizes and over alternative hypotheses, so as to ensure a correct calibration of the test. This point, and more generally the question of Bayesian hypothesis testing and its application to phylogenetic problems, certainly deserve further investigation.

In contrast, cross-validation, being insensitive to the cost mediated by the prior on the effect size, will often incorrectly choose the alternative in this hypothesis testing example. More generally, cross-validation will often fail at suppressing minor but irrelevant fluctuations and redundancies from the output and, as a result, will not be asymptotically consistent in true model identification (Shao, 1993). However, this may be the price to pay, in order to obtain a model selection criterion that is sufficiently flexible in other contexts and for other purposes, such as fitting a sufficiently fine-grained mixture to a complex distribution of random effects. The different aims of model selection, testing hypotheses or finding the best-approximating model, just entail different compromises (Shibata, 1986).

### Implications for Bayesian model averaging

Model averaging is a powerful feature of Bayesian inference, making it possible to consider large combinatorial spaces of model configurations, while integrating uncertainty over models, effect sizes and nuisances (Fragoso *et al*., 2017; Hoeting *et al*., 2000). However, Bayesian model averaging implicitly relies on marginal likelihoods. Therefore, when used in combination with uninformative priors, it will also be biased in favor of the simpler models, just like marginal likelihoods and Bayes factors in the context of explicit model selection. This potentially concerns several previously introduced approaches, implementing model averaging over nucleotide substitution models (Huelsenbeck *et al*., 2004), over the number of components of a mixture (Evans & Sullivan, 2012), or over the number of change points of a non-homogeneous substitution model along the phylogeny (Blanquart & Lartillot, 2006). In the cases just cited, an uninformative prior is used, not just for the global parameters of the model, but also for the replicated items (the exchange rates, the mixture components or the effect sizes associated with each change point). As a result, there is a tendency to over-penalize the more complex model configurations, in spite of the fact that those might be empirically more adequate.

Over-penalization in the context of Bayesian model averaging can be mitigated by the use of a hierarchical prior over the replicated items. For instance, in the case of mixture models, using a hyper-parameterized prior over component-specific parameters will make the model averaging approach less conservative and thus empirically better fitting – as can be seen when comparing the hierarchical (free-hyper) version of the CAT model with its non-hierarchical (fix-hyper) version (Table 2). Similarly, under the change-point model, hyperparameterizing the distribution of the effect sizes upon each transition will result in a more flexible and empirically more adequate model.

All of these points certainly need further exploration and formalization. They also raise a more fundamental question. As mentioned above, marginal likelihoods incorporate a component corresponding to the total cost of fitting, or equivalently, to the total learning work done upon going from the prior to the posterior. The use of hierarchical priors in Bayesian model averaging, by borrowing strength across replicated items, essentially reduces the distance between the prior and the posterior at the level of the replicated items, and thus reduces the cost of fitting. However, it is not clear whether it suppresses this cost entirely. If not, then this suggests that Bayesian model averaging might have a general tendency to be over-penalizing, compared to what could be achieved using more aggressive non-Bayesian model fitting approaches.

### Cross-validation and wAIC: numerical considerations

In contrast to the marginal likelihood, cross-validation appears to be relatively well-behaved, if the aim is to select the most accurate model. However, it requires some care, both for defining the specific details of the CV procedure and for implementing a reliable numerical approach. In this respect, joint k-fold CV gives reasonable results, but it is impractical. There are numerical issues with the naive importance sampling approach, which can lead to a serious underestimation of the CV score, in particular for higher-dimensional models.

The joint k-fold CV approach implemented by naive IS has been used previously for comparing site-heterogeneous and site-homogeneous models (e.g. Philippe *et al*., 2011; Pisani *et al*., 2015; Simion *et al*., 2017). In most cases, site-heterogeneous models have been found as the best fitting models. Importantly, the effective numerical bias of nIS is in favor of less parameter-rich models. This bias, combined with the fact that training has been conducted on datasets of size smaller than the working size, implies that the fit of the site-heterogeneous models has been underestimated thus far. As seen in the case of the elongation-factor dataset (Table 1), the numerical error is the primary cause of the underestimation, although under-training could represent a more substantial contribution in those cases where CV was applied to small subsets of the original data matrix.

The alternative numerical approach used here for computing the joint k-fold CV score, based on sequential importance sampling, is much more accurate than nIS. However it is computationally prohibitive. Of note, there are more sophisticated approaches than the one recruited here for implementing sIS, based on particle filters (Wang *et al*., 2016), which have better Monte Carlo properties than the naive version explored here. However, the single-site importance sampling variance observed here suggests that such particle filters will require many particles and will thus be computationally expensive on data sets of realistic size. More fundamentally, joint k-fold CV does not bring any advantage, compared to its sitewise counterpart, which is numerically much more stable and has a better conceptual justification (see Supplementary Material, section 4).

Leave-one-out cross-validation stands out as the computationally most efficient and most easily implemented cross-validation approach. LOO-CV also has a clear asymptotic connection with information criteria, and more specifically with wAIC. Both can be obtained from a single pass over the MCMC sample, but LOO-CV may require larger MCMC sample sizes (or less thinning) than wAIC to pass the quality checks. The numerical errors appear to be globally well controlled. However, as it stands, the approach is not completely tight. Under small data size, and especially for complex models (here, CAT), there is a non-negligible fraction of sites for which the quality checks for LOO-CV issue a warning about a possible risk of numerical instabilities. Although this did not seem to have any meaningful impact in the present case, nevertheless, these are by nature rare events, which may have been missed in the context of the limited benchmark conducted here and could have a bigger impact in other instances.

One possible work-around for this problem would be to complement LOO-CV and wAIC with sitewise k-fold CV whenever the fraction of sites with low quality checks is high or if a particularly tight evaluation is needed. This approach would present several advantages. Sitewise k-fold CV is more stable than LOO-CV. In addition, LOO-CV is numerically biased in favored of more parameter-rich models, whereas sitewise k-fold CV is biased in favor of less parameter-rich models, both for numerical reasons and because it relies on a smaller training dataset. These opposite biases would make it possible to obtain tighter bounds on the relative fit of any pair of models. With *k* = 5, the impact of under-sized training should be rather moderate, and it would multiply the overall computational cost by a factor at most 6.

In the end, the following practical recommendations are proposed for finding the best approximating model. In the non-problematic cases, LOO-CV and the wAIC represent the most practical and reliable approaches. Both can be obtained from a single pass over the MCMC sample, but LOO-CV may require larger MCMC sample sizes (or less thinning) than wAIC to pass the quality checks. Thus, if the dataset is sufficiently large (*n* > 5000 aligned positions), the wAIC could be used by default, whereas if the dataset is small, then LOO-CV should be preferred. In terms of quality checks, reasonable criteria for good (reasonably good) estimation of the relative fit between two models would be that the mean ESS across sites should be at least 500 (50) for the two models, and that the proportion of sites with low quality measures (ESS < 10 or 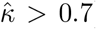) should be at most 5% (10%) for the two models. When those quality requirements are not met, then sitewise k-fold CV could be used as a double-check.

### Practical consequences and perspectives

The arguments exposed here in favor of LOO-CV and wAIC over Bayes factors for model approximation purposes are at odds with the general perception in the applied Bayesian community that Bayes factors represent a general gold standard for model selection (Kass & Raftery, 1995; Lartillot & Philippe, 2006; Xie *et al*., 2011; Oaks *et al*., 2019). This raises the question of the practical consequences of the use of Bayes factors thus far, in situations where cross-validation might have represented a logically more adequate criterion. As illustrated by the analysis of the normal case (Figure 1), for large datasets and for models that don’t differ too much in their dimensionality, all model selection approaches agree in their selection. Thus, in practice, previous results based on the application of Bayes factors on large datasets, such as multi-gene phylogenetic analyses, are unlikely to be qualitatively incorrect, although the case is less clear for smaller-scale analyses. In any case, perhaps a more fundamental contribution of the present analysis is just to facilitate Bayesian model selection, by providing simple guidelines, but also, by making the computational problem of accurately estimating marginal likelihoods practically less relevant.

Still, one main limitation of the methods explored here is that they are valid only in the context of i.i.d. models. For more complex model design, in particular with gene-specific effects in a multi-gene analysis (Suchard *et al*., 2003; Fan *et al*., 2011), or when combining sequence data, fossil information and phenotypic or life-history traits (Lartillot & Poujol, 2011; Zhang *et al*., 2016; Gavryushkina *et al*., 2016), it is less obvious how to define a correct model selection approach and a meaningful asymptotic analysis.

Of note, this limitation also concerns information criteria classically used in a maximum likelihood framework. There have been many illegitimate applications of criteria such as AIC (or BIC) in non i.i.d. settings. Modified versions of these two criteria have been proposed for the specific case of partition models (Seo & Thorne, 2018; Susko & Roger, 2020). It would be useful to develop a wAIC equivalent of the modified AIC that was proposed in this context and, more generally, in the context of other non-iid settings.

Finally, and apart from the practical considerations, the asymptotic theory behind the development of information criteria such as wAIC also suggests an interesting frequentist perspective on Bayesian inference, opening to more general questions, such as efficiently estimating the sampling bias, variance, error, and more general measures of the frequentist risk of the Bayesian estimators, all of which are worth further exploration.

## Supporting information

Supplementary methods, results, figures and table

## Data and software availability

All methods for phylogenetic models were implemented in PhyloBayes (Lartillot *et al*., 2013), version 1.9, available at https://github.com/bayesiancook/pbmpi. The empirical data used here are also available through this repo, along with example scripts. The methods for the analytical and Monte Carlo results under the normal model are available at https://github.com/bayesiancook/normcv.

## Acknowledgements

The Author wishes to thank Marie-Laure Delignette, Philippe Veber, Nicolas Rodrigue and Hervé Philippe for discussions, the Reviewers for their thoughtful comments on the manuscript, and Nicolas Rodrigue for granting access to the computational ressources of his team.

## References

Aho, K., Derryberry, D. & Peterson, T. 2014 Model selection for ecologists: the worldviews of AIC and BIC. Ecology, 95(3), 631–636.

Akaike, H. 1974 A new look at the statistical model identification. IEEE Trans. Automat. Contr., 19(6), 716–723.

Baele, G. & Lemey, P. 2013 Bayesian evolutionary model testing in the phylogenomics era: matching model complexity with computational efficiency. Bioinformatics, 29(16), 1970–1979.

Baele, G., Lemey, P., Bedford, T., Rambaut, A., Suchard, M. A. & Alekseyenko, A. V. 2012a Improving the accuracy of demographic and molecular clock model comparison while accommodating phylogenetic uncertainty. Mol. Biol. Evol., 29(9), 2157–2167.

Baele, G., Lemey, P. & Vansteelandt, S. 2013 Make the most of your samples: Bayes factor estimators for high-dimensional models of sequence evolution. BMC Bioinformatics, 14, 85.

Baele, G., Li, W. L. S., Drummond, A. J., Suchard, M. A. & Lemey, P. 2012b Accurate Model Selection of Relaxed Molecular Clocks in Bayesian Phylogenetics. Mol. Biol. Evol.

Bartlett, M. S. 1957 A comment on D. V. Lindley’s statistical paradox. Biometrika, 44, 533–534.

Berger, J. 2006 The case for objective Bayesian analysis. Bayesian Analysis, 1(3), 385–402.

Berger, J. O. 1985 Statistical Decision Theory and Bayesian Analysis. New-York: Springer-Verlag, 1985th edn.

Bernardo, J. M. & Smith, A. F. M. 1994 Bayesian theory. Chichester, UK: John Wiley & Sons, Inc.

Blanquart, S. & Lartillot, N. 2006 A Bayesian compound stochastic process for modeling nonstationary and nonhomogeneous sequence evolution. Mol. Biol. Evol., 23(11), 2058–2071.

Breiman, L., Friedman, J., Stone, C. J. & Olshen, R. A. 1984 Classification and Regression Trees. Taylor & Francis.

Brown, J. M. & Thomson, R. C. 2017 Bayes Factors Unmask Highly Variable Information Content, Bias, and Extreme Influence in Phylogenomic Analyses. Syst. Biol., 66(4), 517–530.

Bujaki, T. & Rodrigue, N. 2022 Bayesian Cross-Validation Comparison of Amino Acid Replacement Models: Contrasting Profile Mixtures, Pairwise Exchangeabilities, and Gamma-Distributed Rates-Across-Sites. J. Mol. Evol., 90(6), 468–475.

Burnham, K. P. & Anderson, D. R. 2002 Model Selection and Multimodel Inference: a practical information-theoretic approach. New-York: Springer, 2nd edn.

Celeux, G., Forbes, F., Robert, C. P. & Titterington, D. M. 2006 Deviance Information Criteria for Missing Data Models. Bayesian Analysis, 1(4), 651–674.

Chen, M. H., Shao, Q. M. & Ibrahim, J. G. 2012 Monte Carlo Methods in Bayesian Computation. Springer Series in Statistics. Springer New York.

Efron, B. 1986 How Biased is the Apparent Error Rate of a Prediction Rule? Journal of the American Statistical Association, 81(394), 461–470.

Evans, J. & Sullivan, J. 2012 Generalized mixture models for molecular phylogenetic estimation. Syst. Biol., 61(1), 12–21.

Fan, Y., Wu, R., Chen, M.-H., Kuo, L. & Lewis, P. O. 2011 Choosing among partition models in Bayesian phylogenetics. Mol. Biol. Evol., 28(1), 523–532.

Fragoso, T. M., Bertoli, W. & Neto, F. L. 2017 Bayesian model averaging: A systematic review and conceptual. International Statistical Review, 86(1), 1–28.

Gavryushkina, A., Heath, T. A., Ksepka, D. T., Stadler, T., Welch, D. & Drummond, A. J. 2016 Bayesian Total-Evidence Dating Reveals the Recent Crown Radiation of Penguins. Syst. Biol.

Geisser, S. 1975 The predictive sample reuse method with application. Journal of the American Statistical Association, 70, 320–328.

Geisser, S. & Eddy, W. 1979 A predictive approach to model selection. Journal of the American Statistical Association, 74, 153–160.

Gelfand, A. E. 1996 Model determination using sampling-based methods. In Markov chain monte carlo in practice (eds W. R. Gilks, S. Richardson & D. J. Spiegelhalter), pp. 145–162. Chapman & Hall/CRC.

Gelfand, A. E., Dey, D. K. & Chang, H. 1992 Model determination using predictive distributions with implementation via sampling-based methods. In Bayesian statistic, 4th *edn* (eds J. M. Bernardo, J. O. Berger, A. P. Dawid & A. F. M. Smith), pp. 147–167. Oxford: Oxford University Press.

Gelman, A., Hwang, J. & Vehtari, A. 2014 Understanding predictive information criteria for Bayesian models. Stat Comput, 24, 997–1016.

Goldman, N. 1993 Statistical tests of models of DNA substitution. J. Mol. Evol., 36(2), 182–198.

Hoeting, J. A., Madigan, D., Raftery, A. E. & Volinsky, C. T. 2000 Bayesian Model Averaging: A Tutorial. Statistical Science, 14(4), 382–417.

Huelsenbeck, J. P., Larget, B. & Alfaro, M. E. 2004 Bayesian phylogenetic model selection using reversible jump Markov chain Monte Carlo. Mol. Biol. Evol., 21(6), 1123–1133.

Jeffreys, H. 1935 Some tests of significance, treated by the theory of probability. Proc. Camb. Phil. Soc., 31, 203–222.

Jeffreys, H. 1967 Theory of probability. London: Oxford University Press.

Jones, D. T., Taylor, W. R. & Thornton, J. M. 1992 The rapid generation of mutation data matrices from protein sequences. Comput Appl Biosci, 8(3), 275–282.

Kass, R. E. & Raftery, A. E. 1995 Bayes Factors. Journal of the American Statistical Association, 90(430), 773–795.

Konishi, S. & Kitagawa, G. 1996 Generalised information criteria in model selection. Biometrika, 83(4), 875–890.

Konishi, S. & Kitagawa, G. 2007 Information Criteria and Statistical Modeling. Springer New York.

Kosakovsky Pond, S. L. & Frost, S. D. W. 2005 Not so different after all: a comparison of methods for detecting amino acid sites under selection. Mol. Biol. Evol., 22(5), 1208–1222.

Lartillot, N., Brinkmann, H. & Philippe, H. 2007 Suppression of long-branch attraction artefacts in the animal phylogeny using a site-heterogeneous model. BMC Evol. Biol., 7 Suppl 1, S4.

Lartillot, N., Lepage, T. & Blanquart, S. 2009 PhyloBayes 3: a Bayesian software package for phylogenetic reconstruction and molecular dating. Bioinformatics, 25(17), 2286–2288.

Lartillot, N. & Philippe, H. 2004 A Bayesian mixture model for across-site heterogeneities in the amino-acid replacement process. Mol. Biol. Evol., 21(6), 1095–1109.

Lartillot, N. & Philippe, H. 2006 Computing Bayes factors using thermodynamic integration. Syst. Biol., 55(2), 195–207.

Lartillot, N. & Philippe, H. 2008 Improvement of molecular phylogenetic inference and the phylogeny of Bilateria. Philos. Trans. R. Soc. Lond. B Biol. Sci., 363(1496), 1463–1472.

Lartillot, N. & Poujol, R. 2011 A phylogenetic model for investigating correlated evolution of substitution rates and continuous phenotypic characters. Mol. Biol. Evol., 28(1), 729–744.

Lartillot, N., Rodrigue, N., Stubbs, D. & Richer, J. 2013 PhyloBayes MPI: phylogenetic reconstruction with infinite mixtures of profiles in a parallel environment. Syst. Biol., 62(4), 611–615.

Le, S. Q. & Gascuel, O. 2008 An improved general amino acid replacement matrix. Mol. Biol. Evol., 25(7), 1307–1320.

Lewis, P. O., Xie, W., Chen, M.-H., Fan, Y. & Kuo, L. 2014 Posterior predictive Bayesian phylogenetic model selection. Syst. Biol., 63(3), 309–321.

Lindley, D. V. 1957 A statistical paradox. Biometrika, 44, 187–192.

Nielsen, R. & Yang, Z. 1998 Likelihood models for detecting positively selected amino acid sites and applications to the HIV-1 envelope gene. Genetics.

Oaks, J. R., Cobb, K. A., Minin, V. A. & Leaché, A. D. 2019 Marginal Likelihoods in Phylogenetics: A Review of Methods and Applications. Syst. Biol., 68(5), 681–697.

Pagel, M. & Meade, A. 2004 A phylogenetic mixture model for detecting pattern-heterogeneity in gene sequence or character-state data. Syst. Biol., 53(4), 571–581.

Philippe, H., Brinkmann, H., Copley, R. R., Moroz, L. L., Nakano, H., Poustka, A. J., Wallberg, A., Peterson, K. J. & Telford, M. J. 2011 Acoelomorph flatworms are deuterostomes related to Xenoturbella. Nature, 470(7333), 255–258.

Philippe, H., Lartillot, N. & Brinkmann, H. 2005 Multigene analyses of bilaterian animals corroborate the monophyly of Ecdysozoa, Lophotrochozoa, and Protostomia. Mol. Biol. Evol., 22(5), 1246–1253.

Pisani, D., Pett, W., Dohrmann, M., Feuda, R., Rota-Stabelli, O., Philippe, H., Lartillot, N. & Wörheide, G. 2015 Genomic data do not support comb jellies as the sister group to all other animals. Proceedings of the National Academy of Sciences, 112(50), 15 402–15 407.

Plummer, M. 2008 Penalized loss functions for Bayesian model comparison. Biostatistics, 9(3), 523–539.

Raftery, A. E., Newton, M. A., Satagopan, J. M. & Krivitsky, P. N. 2007 Estimating the Integrated Likelihood via Posterior Simulation Using the Harmonic Mean Identity. Bayesian Statistics, 8, 1–45.

Ronquist, F., Kudlicka, J., Senderov, V., Borgström, J., Lartillot, N., Lundén, D., Murray, L., Schön, T. B. & Broman, D. 2021 Universal probabilistic programming offers a powerful approach to statistical phylogenetics. Communications Biology, pp. 1–10.

Schrempf, D., Lartillot, N. & Szöllősi, G. 2020 Scalable empirical mixture models that account for across-site compositional heterogeneity. Mol. Biol. Evol.

Schwarz, G. 2006 Estimating the Dimension of a Model. Ann. Statist., 6(2), 461–464.

Seo, T.-K. & Thorne, J. L. 2018 Information Criteria for Comparing Partition Schemes. Syst. Biol., 67(4), 616–632.

Shao, J. 1993 Linear Model Selection by Cross-Validation. Journal of the American Statistical Association, 88(422), 486–494.

Shibata, R. 1986 Consistency of Model Selection and Parameter Estimation. Journal of applied probability, 23, 127–141.

Shibata, R. 1989 Statistical aspects of model selection. In From data to model (ed. J. C. Willems), pp. 215–240. Springer New York.

Shimodaira, H. 2004 Approximately unbiased tests of regions using multistep-multiscale bootstrap resampling. Ann. Statist., 32(6), 2616–2641.

Simion, P., Philippe, H., Baurain, D., Jager, M., Richter, D. J., Di Franco, A., Roure, B., Satoh, N., Quéinnec, E. et al. 2017 A Large and Consistent Phylogenomic Dataset Supports Sponges as the Sister Group to All Other Animals. Curr. Biol., 27(7), 958–967.

Smyth, P. 2000 Model selection for probabilistic clustering using cross-validated likelihood - Springer. Stat Comput.

Spiegelhalter, D. J., Best, N. G. & Carlin, B. P. 2002 Bayesian measures of model complexity and fit. J. R. Statist. Soc. B, 64, 583–639.

Spiegelhalter, D. J., Best, N. G., Carlin, B. P. & Van der Linde, A. 2014 The deviance information criterion: 12 years on. J. R. Statist. Soc. B, 76(3), 485–493.

Stone, M. 1974 Cross-validatory choice and assessment of statistical predictions. J. R. Statist. Soc. B, pp. 111–147.

Stone, M. 1977 An asymptotic equivalence of choice of model by cross-validation and Akaike’s criterion. J. R. Statist. Soc. B, pp. 44–47.

Suchard, M. A., Kitchen, C. M. R., Sinsheimer, J. S. & Weiss, R. E. 2003 Hierarchical Phylogenetic Models for Analyzing Multipartite Sequence Data. Syst. Biol., 52(5), 649–664.

Suchard, M. A., Weiss, R. E. & Sinsheimer, J. S. 2001 Bayesian selection of continuous-time Markov chain evolutionary models. Mol. Biol. Evol., 18(6), 1001–1013.

Sullivan, J. & Joyce, P. 2005 Model selection in phylogenetics. Annu. Rev. Ecol. Evol. Syst., pp. 445–466.

Susko, E., Lincker, L. & Roger, A. J. 2018 Accelerated Estimation of Frequency Classes in Site-Heterogeneous Profile Mixture Models. Mol. Biol. Evol., 35(5), 1266–1283.

Susko, E. & Roger, A. J. 2020 On the Use of Information Criteria for Model Selection in Phylogenetics. Mol. Biol. Evol., 37(2), 549–562.

Thomas, V., Pedregosa, F., van Merriënboer, B., Mangazol, P.-A., Bengio, Y. & Le Roux, N. 2020 On the interplay between noise and curvature and its effect on optimization and generalization. In Proceedings of the 23rdinternational conference on artificial intelligence and statistics (aistats).

Vehtari, A., Gelman, A. & Gabry, J. 2016 Practical Bayesian model evaluation using leave-one-out cross-validation and WAIC. Stat Comput, 27(5), 1413–1432.

Vrieze, S. I. 2012 Model selection and psychological theory: a discussion of the differences between the Akaike information criterion (AIC) and the Bayesian information criterion (BIC). Psychol Methods, 17(2), 228–243.

Wang, L., Bouchard-Côté, A. & Doucet, A. 2016 Bayesian Phylogenetic Inference Using a Combinatorial Sequential Monte Carlo Method. Journal of the American Statistical Association, 110(512), 1362–1374.

Watanabe, S. 2001 Algebraic geometrical methods for hierarchical learning machines. Neural Netw, 14(8), 1049–1060.

Watanabe, S. 2007 Almost All Learning Machines are Singular. IEEE Symposium on Foundations of Computational Intelligence.

Watanabe, S. 2009 Algebraic Geometry and Statistical Learning Theory. Cambridge Monographs on Applied and Computational Mathematics. Cambridge University Press.

Watanabe, S. 2010a Asymptotic Equivalence of Bayes Cross Validation and Widely Applicable Information Criterion in Singular Learning Theory. The Journal of Machine Learning Research, 11, 3571–3594.

Watanabe, S. 2010b Equations of states in singular statistical estimation. Neural Netw, 23(1), 20–34.

Whelan, S. & Goldman, N. 2001 A general empirical model of protein evolution derived from multiple protein families using a maximum-likelihood approach. Mol. Biol. Evol., 18(5), 691–699.

Xie, W., Lewis, P. O., Fan, Y., Kuo, L. & Chen, M.-H. 2011 Improving marginal likelihood estimation for Bayesian phylogenetic model selection. Syst. Biol., 60(2), 150–160.

Zhang, C., Stadler, T., Klopfstein, S., Heath, T. A. & Ronquist, F. 2016 Total-Evidence Dating under the Fossilized Birth-Death Process. Syst. Biol., 65(2), 228–249.

Zhang, J., Nielsen, R. & Yang, Z. 2005 Evaluation of an improved branch-site likelihood method for detecting positive selection at the molecular level. Mol. Biol. Evol.

Zhang, P. 1993 Model selection via multifold cross validation. Tha Annals of Statistics, 21(1), 299–313.

