## Supplementary methods, results, figures and table for "Identifying the best approximating model in Bayesian phylogenetics: Bayes factors, cross-validation or wAIC?"

Nicolas Lartillot

*Laboratoire de Biométrie et Biologie Evolutive, UMR CNRS 5558, Université Lyon 1, Villeurbanne, France.*

**Running head:** Bayes factors, Cross-validation and wAIC

cross-validation, Bayes factor, marginal likelihood, model comparison, wAIC

### 1 Additional methods on Monte Carlo

The notations are briefly recalled. The dataset is noted $X = (X_i)_{i=1..n}$. For notational convenience, it is assumed that, for cross validation, the first $q$ data points were used for training, and the last $r$ for validation, with $q$ and $r$ depending on the exact cross-validation approach. $X_{a:b}$ denotes the set of observations $(X_i)_{a \leq i \leq b}$. When $q = 0$, $X_{1:q}$ is the empty set. The index $i = 1..n$ runs over data points, and $t = 1..T$ over the parameter configurations sampled by MCMC.

#### 1.1 Estimating Monte Carlo effective sample sizes

All approaches introduced above, nIS, CPO and sIS, are based on Monte Carlo estimators in which an expectation of some function $W(\theta)$ under some distribution of density $\pi(\theta)$ is replaced by an average over $T$ parameter configurations sampled from this distribution:

$$\int W(\theta)\pi(\theta)d\theta \simeq \frac{1}{T}\sum_{t=1}^{T} W(\theta_t), \tag{1}$$

where $\theta_t \sim \pi(\theta)$, for $t = 1..T$. The quality of this Monte Carlo estimate depends on the variance of $W(\theta)$ under the distribution defined by $\pi(\theta)$. A useful statistic in this respect is the effective sample size (ESS). The ESS is obtained by first defining the normalized weights:

$$w_t = \frac{W(\theta_t)}{\sum_{t'=1}^{T} W(\theta_{t'})} \tag{2}$$

and then computing:

$$\texttt{ESS} = \left(\sum_{t=1}^{T} w_t^2\right)^{-1} \tag{3}$$

If $W(\theta)$ has a small variance, then $w_t \simeq 1/T$ for all $T$ and the ESS is essentially equal to $T$. Conversely, when $W(\theta)$ has a large variance, then one single sample tends to dominate, such that all $w_t$'s are close to 0, except for one of them which is close to 1. In that case, the ESS is essentially equal to 1. The application of this general idea to each of the methods described above is straighforward:

- nIS for joint k-fold CV: a single ESS is computed, using $W(\theta) = p(X_{q+1:n} \mid \theta)$;
- nIS for site-wise k-fold CV and sIS: an ESS is computed for each data point, using $W_i(\theta) = p(X_i \mid \theta)$ for point $i$;
- CPO: an ESS is computed for each data point, using $W_i(\theta) = p(X_i \mid \theta)^{-1}$ for point $i$;

For all methods that return an ESS for each data point, the mean ESS over all data points is reported, along with the fraction of data points for which the ESS is less than 10. This last statistic gives an idea of how much of the score that is eventually reported is contributed by terms for which the Monte Carlo error is potentially not well controlled.

#### 1.2 Automatic tuning of the Monte Carlo on a site-specific basis for sIS

As mentioned in the main text, the quality of the estimate of site-specific predictive score used by sIS, i.e. $p(X_i \mid X_{1:i-1})$, depends on the variance of the log-likelihood $\ln p(X_i \mid \theta)$ under the partial posterior $p(\theta \mid X_{1:i-1})$, which itself varies greatly between data points. Accordingly, the number of Monte Carlo samples is tuned separately for each site, by first running a small number of cycles at step $i$ to estimate the variance of the log-likelihood and then proceed with a number of cycles $T_i$ determined based on this variance estimate.

A heuristic argument for deciding how $T_i$ should scale as a function of $v_i$ is as follows. If the $L_{it}$'s were normally distributed, of variance $v_i$, then the variance of $\ln L_i$, i.e. the log-transformed Monte Carlo estimate given by equation **??** would scale as $V \sim \frac{e^{v_i}}{T_i}$. Inverting this equation, this suggests that, in order to target a given fixed variance for all data points, $T_i$ should scale as $e^{v_i}$. Taken together, these observations lead to a second version of the algorithm, which runs a follows. Starting from a parameter configuration sampled from the prior $\theta_0 \sim p(\theta)$, at step $i = 1..n$:

- the MCMC is run for B cycles, giving B new parameter configurations $\theta_{it}$, for $t = 1..B$, approximately under the partial posterior distribution $p(\theta \mid X_{1:i-1})$. For each $t = 1..B$, the likelihood of the next data point is calculated, i.e. $L_{it} = p(X_i \mid \theta_{it})$.

- the sample variance $\hat{v}_i$ of the $\ln L_{it}$'s is computed and used to determine $T_i$, using the following rule: $T_i = \min(T_0 e^{\hat{v}_i - v_0}, T_{max})$, which thus implements an exponential scaling of $T_i$ as a function of $v_i$, targeting a sample size of $T_0$ for sites for which have a variance equal to $v_0$, and truncated at $T_{max}$.

- the MCMC is run for another series of $T_i$ cycles, yielding a new sample of parameter configurations $\theta'_{it}$, for $t = 1..T_i$; the likelihood factors for the next data point are computed, $L'_{it} = p(X_i \mid \theta'_{it})$, for $t = 1..T_i$.

- this second series of likelihoods is used to compute the arithmetic average $L_i$

Of note, the preliminary run of $B$ cycles at the beginning of each step also contributes to equilibrating the MCMC just after the addition of the last data point. In some cases, there is still a small minority of data points for which $v_i$ may be too large, such that $T_i = T_{max}$ and the Monte Carlo error may not be well controlled. On the other hand, if they represent a small fraction of the data, their contribution to the total error should be small. This point is checked in a second step, based on the effective sample size (such as defined above, subsection 1.1).

##### 1.3 Reference distributions for sIS

The general idea for implementing the empirical reference distributions for sIS under phylogenetic models is to use, for each parameter, the same parametric distributions as the original prior, such as implemented in the model, and to modify the value of its hyper-parameters so as to match as closely as possible the mean and the variance of the posterior distribution, such as estimated based on preliminary run (moment-matching).

The parameters for which this is done are: the branch lengths ($l_t$, $t = 1..2P-3$, where $T$ is the number of taxa), the shape parameter $\alpha$ of rates across sites, the equilibrium frequencies over amino-acids under JTT, LG and GTR ($\pi_a$, for $a = 1..20$), the relative exchange rates under GTR ($\rho_{ab}$, for $1 \leq a < b \leq 20$), and the hyper-parameters $\kappa$, $\beta$ and $\pi_0$ for the mixture of the CAT model. The priors implemented in PhyloBayes for those parameters are first recalled:

- branch lengths: $l_j \sim \texttt{Gamma}(1, \mu)$ (exponential of mean $\mu^{-1}$);
- branch length hyper-parameter $\mu \sim \texttt{Gamma}(1, 0.1)$ (exponential of mean 10);
- shape parameter of rates across sites: $\alpha \sim \texttt{Gamma}(1, 1)$ (exponential of mean 1);
- for JTT, LG, GTR: equilibrium frequencies over amino-acids $\pi \sim \texttt{Dirichlet}(1, 1, ...)$
- for GTR, relative exchangeabilities: $\rho_{it} \sim \texttt{Gamma}(1, 1)$ (exponential or mean 1; equivalently, the renormalized exchange rates are from a uniform Dirichlet distribution over the simplex);

for the CAT model:

- mixture weights are from a stick-breaking process (truncated at 1000) of parameter $\kappa \sim \texttt{Gamma}(1, 0.1)$ (exponential of mean 10);
- mixture profiles are i.i.d. from base distribution: $\pi_k \sim \texttt{Dirichlet}(\beta\pi_0)$;

- for the fixhyper version of the CAT model, $\beta = 20$ and $\pi_0 = (0.05, 0.05, ...)$ (as a result, the $\pi_k$'s are i.i.d. from a uniform base distribution)

- for free hyper: $\beta$ and $\pi_0$ themselves hyperparameters with priors: $\beta \sim \texttt{Gamma}(1, 0.05)$ (exponential of mean 20); and $\pi_0 \sim \texttt{Dirichlet}(1, 1, ...)$.

Thus, for some specific parameter of the model generically noted $\theta$, the sample mean and variance (noted $\hat{M}[\theta]$ and $\hat{V}[\theta]$), are first computed over the preliminary MCMC run. Next, these sample means and variances are converted by moment matching into shape and rate parameters of gamma distributions, or into concentration parameters of Dirichlet distributions, depending on the parameter under focus. Finally, these empirical hyperparameters are used to design the family of reference priors. More specifically, when $\theta$ is gamma distributed of shape $a_0$ and rate $b_0$ under the original prior (e.g. brancg lengths, $\alpha$, relative exchange rates), then the empirical shape and scale parameters are defined as $\hat{a}(\theta) = \hat{M}[\theta]^2/\hat{V}[\theta]$ and $\hat{b}(\theta) = \hat{M}[\theta]/\hat{V}[\theta]$, and the reference distribution, corresponding to the density $p_\epsilon(\theta)$ introduced above (Variance reduction subsection), is then defined as:

$$\theta \sim \texttt{Gamma}(\epsilon\, a_0 \,+\, (1-\epsilon)\,\hat{a}(\theta), \epsilon b_0 \,+\, (1-\epsilon)\,\hat{b}(\theta)).$$

When $\theta = (\theta_m)_{m=1..M}$ is a frequency vector (such as equilibrium frequencies over amino-acids), from a Dirichlet of concentration $(a_m^0)_{m=1..M}$ under the original prior, the empirical concentration parameters are defined as $\hat{a}_m = \hat{C}\hat{M}[\theta_m]$, where

$$\hat{C} \;=\; \left(\sum_m \hat{M}[\theta_m](1-\hat{M}[\theta_m])\right) / \left(\sum_m \hat{V}[\theta_m]\right) - 1.$$

The reference distribution is then defined as:

$$\theta \sim \texttt{Dirichlet}(\epsilon\, a_1^0 \,+\, (1-\epsilon)\,\hat{a}_1(\theta), \ldots, \epsilon\, a_M^0 \,+\, (1-\epsilon)\,\hat{a}_M(\theta)).$$

With these definitions, when $\epsilon = 0$, the reference distribution has a mean and a variance equal to the posterior estimates obtained under the preliminary run, while it reduces to the original prior distribution when $\epsilon = 1$.

##### 1.4 Estimation of the bias for nIS, sIS and CPO

The general method for estimating the bias in the case of nIS, sIS and CPO is presented. This method proceeds from the same idea in all three cases, although with some differences in the details, depending on the specific MCMC approach. For all three methods, a set of $M$ independent Monte Carlo runs is first conducted (on the same data replicate, i.e. under the same permutation or the data points, in the case of cross-validation). A given run of the Monte Carlo algorithm produces, for each data point, a vector of $T$ likelihood scores. These arrays are generically noted $L_{it}^m$, for site $i = 1..n$, for step $t = 1..T$ and run $m = 1..M$. These likelihood scores are then combined differently, depending on the method, such as described in the last subsection.

To set up a general formalism valid for all methods, in the following, the quantity to be estimated by each of these methods is generically noted $z_i$ at step $i = 1..n$. Thus, for site-wise k-fold CV (nIS):

$$z_i = p(X_i \mid X_{1:q}),\ i = q+1..n$$

for joint k-fold CV and marginal likelihood (sIS):

$$z_i = p(X_i \mid X_{1:i-1}),\ i = q+1..n$$

and for LOO-CV (CPO):

$$z_i = p(X_i \mid X_{(i)})^{-1},\ i = 1..n$$

With these definitions, $z$ always corresponds to the quantity for which the corresponding Monte Carlo approach will give an unbiased estimate on the natural scale. As described in the last subsection, for the contribution for site $i$ and over run $m$, this unbiased estimator, noted $\hat{z}_i^m$, is the arithmetic mean over the Monte Carlo samples of the observation-specific likelihoods for sitewise k-fold CV (nIS), joint k-fold CV and the marginal likelihood (sIS):

$$\hat{z}_i^m = \frac{1}{T} \sum_{t=1}^{T} L_{it}^m$$

and the arithmetic mean of their inverse for LOO-CV (CPO):

$$\hat{z}_i^m = \frac{1}{T} \sum_{t=1}^{T} 1/L_{it}^m$$

These estimates are then log-transformed, giving $\hat{y}_i^l = \ln \hat{z}_i^m$ for nIS and sIS, and $\hat{y}_i^m = -\ln \hat{z}_i^m$ for CPO, to account for the fact that $z_i$ is the inverse of the LOO-CV score. For all methods, the global per-site score, generically noted $\hat{y}^m$ for run $m$, is then the mean over data points of the validation set of the log-transformed estimates:

$$\hat{y}^m = \frac{1}{r} \sum_{i=q+1}^{n} \hat{y}_i^m \tag{4}$$

and finally, these $M$ independent estimates are averaged over all runs:

$$\hat{y} = \frac{1}{M}\sum_m \ln \hat{y}^m$$

giving a first raw estimate.

As mentioned above, $\hat{z}_i^m$ is unbiased as an estimator of $z_i$. However, because of the log-transformation, $\hat{y}_i^m$, and thus also $\hat{y}$, are biased. This bias can be estimated using a quadratic expansion of the logarithm. The general idea is that, for any random variable $Z$ with expectation $z$, up to quadratic terms:

$$Y = \ln Z = \ln((Z-z)+z) = \ln z + \frac{Z-z}{z} - \frac{(Z-z)^2}{2z^2} + o((Z-z)^2)$$

and thus

$$\begin{aligned} E[Y] &\simeq \ln z + 0 - \frac{var[Z]}{2E[Z]^2} \\ &\simeq \ln z - \frac{1}{2} var[Y] \end{aligned}$$

In other words, if $Z$ is an unbiased estimator of $z$, then $Y = \ln Z$, as an estimator of $\ln z$, has a negative bias approximately equal to half of its own variance. For CPO, $Y = -\ln Z$ and thus the bias is positive but has otherwise the same expression. This development will be valid if the variance of $Y$ is small. In the present case, one can rely on the fact that the estimator given by equation 4 is an average of the individual contributions given by each data point of the validation set, each of which has a small variance. Accordingly, the bias should be estimated separately for each $\hat{y}_i$, and then the total bias can be computed by averaging over data points.

Specifically, an estimate of the bias $\hat{b}$ of the estimator $\hat{y}$ is obtained by first computing the mean and the variance of the site-specific scores over runs. Thus, for site $i = 1..n$:

$$\begin{aligned} \bar{y}_i &= \frac{1}{M}\sum_m y_i^m \\ \hat{v}_i &= \frac{1}{M-1}\sum_m (y_i^m - \bar{y}_i)^2 \end{aligned}$$

Note that the unbiased estimator of the variance is used here. Assuming that these site-specific variances are small, the estimator of the bias introduced above can be used, i.e. $\hat{b}_i = -0.5\,\hat{v}_i$ for nIS and sIS, and $\hat{b}_i = 0.5\,\hat{v}_i$ for CPO. The overall bias is thus:

$$\hat{b} = \frac{1}{r}\sum_i \hat{b}_i \tag{5}$$

giving the final debiased estimate $\tilde{y} = \hat{y} - \hat{b}$.

Of note, the site-specific sample variances (the $\hat{v}_i$'s) are potentially noisy, as estimators of the corresponding true variances. However, they are unbiased estimators of the true Monte Carlo variances. Similarly, the overall bias, which is a linear function of the $v_i$'s, is thus also unbiased. Provided that the $\hat{v}_i$'s are not too strongly correlated across sites, their average over site entailed by equation 5 contributes a lot to stabilizing the bias estimate given by equation 5. As a result, in practice, only $M = 2$ runs are enough to give a good estimate of the bias, and running more replicates (e.g. $M = 10$) does not yield notable improvement. The number of Monte Carlo samples per data point, $T$, on the other hand, should be large enough so as to guarantee that the $\hat{v}_i$'s are all small, in order for the linear estimator of the bias to be valid. In practice, it was found that, in order to obtain a reliable numerical estimate of the bias, the ESS associated to the computation of each $\hat{y}_i^m$ should be greater than 10.

#### 2 General settings across all experiments

For the experiments shown in Table 1, under the normal model, 100 datasets of size $n = 1000$ observations were simulated under the following parameter values: dimension $p = 100$, 300, 1000, true mean $\theta_* = 0.04$, and variance parameter $\sigma^2 = 1$. For each replicate, the analytical values of the log Bayes factor and the relative cross-validation score between the two models were computed (see below, section Analytical results under the normal model). In the case of cross-validation, since the results are ultimately averaged over the 100 independent simulation replicates, only one choice for the splitting of the dataset into a validation and a training set is considered for each replicate, taking the first 800 data points for training and the last 200 for validation. For sIS, the first approach (fixed $T$ for all data points) was used, with $T$ such as indicated in Table 1.

In the case of EF2 (Table 2), for k-fold CV (both joint and site-wise), 10 random reshufflings of the sites of the original dataset were created. Each of these reshuffled version of the dataset was further split into a training and a validation set, on which nIS was applied, running the MCMC for 1100 samples (saving every cycle for JTT, LG, GTR and every 3 cycles for CAT) on the training set and then computing the CV score of the validation set based on the 1000 last samples of the MCMC. For sIS, a preliminary run on the original dataset and under each model was conducted for 1100 samples (saving every 3 cycles for CAT). Empirical posterior means and variances for defining the reference priors for sIS were obtained based on the last 1000 samples of this preliminary run. The self-tuned version of the sIS method was then used, with the following parameter values: $B = 10$,

$P_0 = 30$, $P_{max} = 1000$ and $v_0 = 0.1$, with $M = 2$ independent runs for each data replicate (using the original dataset for computing the marginal likelihood and the 10 random reshufflings of the sites for k-fold CV). For LOO-CV, $M = 2$ independent runs were conducted on the complete dataset and under each model, again for 1100 samples and saving every 3 cycles for CAT, discarding the first 100 samples of burn-in and applying the CPO method on the 1000 remaining samples.

For the experiments shown in Figure 2, the Metazoan dataset was first filtered to remove sites with more than 20% of missing data, leaving a total of 9804 sites. Then, 4 jackknife replicates of sizes $n = 200$ to 800 were randomly sampled. For figure 3, a standard MCMC was run under the complete dataset (9804 sites) and under the LG model, for a total of 1100 samples. Then, 10 posterior predictive replicates were simulated, based on 10 samples regularly spaced across this MCMC chain, discarding the first 100 samples and taking one every 100 samples. Each of these simulated datasets was then jackknifed, yielding replicates of sizes $n = 200$ to $n = 800$. Both the empirical and the simulated jackknife replicates were then used for assessing the fit of the model by marginal likelihood or LOO-CV, using the same settings as for EF2.

For Figure 4, the complete dataset was used, and 4 jackknife replicates of size ranging from $n = 200$ to $n = 16000$ were randomly sampled. The LOO-CV scores and the wAIC were computed by running $L = 2$ independent chains of 1100 samples (saving every 3 cycles for CAT) on each jackknife replicate, using the last 1000 samples, with or without a 10-fold thinning (in which case the Monte Carlo estimates of the posterior averages are based on 100 samples).

### 3 Numerical stability and accuracy of LOO-CV and the wAIC

In this section, the numerical stability and accuracy of Monte Carlo estimation for LOO-CV and wAIC are explored in the case of LG, GTR and CAT-Poisson (CAT-GTR was not considered for computational reasons), using a combination of several approaches.

#### 3.1 Methods

First, several statistics were monitored: the estimated Monte Carlo bias and standard deviation, the mean effective sample size (mean ESS) across sites and the fraction of sites for which the ESS is less than 10. The mean ESS is directly related to the Monte Carlo variance of the estimator. The fraction of sites having a critically low ESS, on the other hand, is an indicator of the risk of not correctly estimating the bias. Two alternative settings were explored for Monte Carlo sample size, $T = 1000$ and $T = 100$, the latter being obtained by thinning the first sample, i.e. taking 1 every 10 samples.

Second, the Pareto-smoothed importance sampling approach for LOO-CV (hereafter called PS-LOO-CV) (Vehtari *et al.*, 2016) was used. The rationale of PS-LOO-CV is to stabilize the harmonic mean estimator entailed by the CPO approach, which may be sensitive to one or a few of the largest importance weights (here, the inverse of the site-specific likelihood values), by fitting a generalized Pareto distribution to the right tail of the empirical series of importance weights (20% largest importance ratios), and this, independently for each data point. The contribution of these 20% importance weights to the harmonic mean is then replaced by the expectation under the generalized Pareto distribution. The method gives, as a by-product, an estimate $\hat{\kappa}$ of the parameter of the generalized Pareto distribution (independently for each site). The higher the value of $\kappa$, the heavier the tails, and as a result, the higher the risk of numerical instabilities. A criterion used here is the fraction of sites for which $\hat{\kappa} > 0.7$.

Third, the three estimators, LOO-CV, PS-LOO-CV and the wAIC, were checked against yet another alternative estimation methods, which in the following is called 2-step CV. This 2-step method method runs as follows: for each replicate (say $X$) on wich LOO-CV and the wAIC were both applied, the model was trained on an independent dataset ($Y$) randomly sampled from the same original phylogenetic dataset, before being validated on $X$. The training and validation sets are both of same size and are non-overlapping. For training, the MCMC was run under the same conditions as for LOO-CV and for the wAIC. For validation, the score is computed sitewise.

More formally, if $X = (X_i)_{i=1..n}$ is the validation dataset and $Y = (Y_i)_{i=1..n}$ the training set, the score returned by 2-step CV is a Monte Carlo esimator of:

$$cv_2 \quad = \quad \frac{1}{n}\sum_{i=1}^{n} \ln p(X_i \mid Y) \tag{6}$$

Estimation runs as for sitewise k-fold CV. That is, for $t = 1..T$, compute the likelihood separately for each data point of the validation data, $L_{it} = p(X_i \mid \theta_t)$, for $i = 1..n$. In a second step, for all $i = 1..n$, compute the arithmetic mean of the $L_{it}$'s over the $T$ Monte Carlo samples, log-transform, and finally, sum all individual contributions across the $n$ data points of the validation set.

With this definition, the 2-step approach provides an estimate of $C(n, 1)$ in the notation of the main article. As a reminder, LOO-CV is an estimate of $C(n-1, 1)$ (see Section 'The theoretical targets of Bayesian cross-validation and the wAIC'). In terms of Monte Carlo errors, LOO-CV has a positive bias, whereas 2-step CV has a negative bias. In both cases, the bias is more pronounced for more parameter-rich models. Thus, when computing the fit of a complex model (such as CAT-Poisson) relative to a simpler model (such as LG), the overall bias is still positive for LOO-CV and negative for 2-step CV. Up to the sampling and the Monte Carlo variance, and up to a small deviation of $1/n$ between there respective targets, the raw (i.e. not de-biased) estimates of LOO-CV and the two-step approach should therefore be on either side of the true predictive fit. Of note, when compared with LOO-CV, the 2-step approach entails an additional sampling variance, which is minimized here by matching the validation set of 2-step CV with the dataset on which LOO-CV is computed, although there still remains an additional component of variance contributed by the choice of the independent training dataset. For the comparison with the wAIC, the expectations are less straightforward: like the 2-step approach, the wAIC has a negative bias. However, it also entails an additional error due to the fact that it is an asymptotic approximation of $C(n, 1)$.

##### 3.2 Results

The Monte Carlo statistics for LOO-CV and wAIC and PS-LOO-CV are summarized in Supplementary Table 1, along with the mean point LOO-CV and wAIC estimates. First, not surprisingly, estimation quality improves with Monte Carlo sample size, and this, based on all measures of estimation quality. Second, in some situations, in particular for small sample size, small data size, and more specifically for the CAT-Poisson model, there is a fraction of sites showing either an ESS $< 10$ or a value of $\hat{\kappa} > 0.7$. The fraction of sites with critically low ESS or critically high value of $\hat{\kappa}$ appears to be the most sensitive criterion for assessing numerical stability: for instance, whenever the estimated fit shows a difference greater than 0.01 unit per site between the two alternative thinning schemes, the fraction of sites with a critically low ESS under the smaller MCMC sample is always above 5%, and conversely. Both estimators, LOO-CV and wAIC, tend to be numerically more stable for larger data size, although the wAIC appears to be numerically more stable than LOO-CV.

Supplementary Figure 1 below gives a graphical account of the estimated fit, according to PS-LOO-CV, LOO-CV, the wAIC and 2-step CV, as a function of datasize, for GTR and CAT-Poisson, relative to LG. Overall, the differences between LOO-CV, the wAIC and 2-step CV are small compared to the difference between models and to the sampling variance. The opposite bias of LOO-CV and 2-step CV is visible, in particular for CAT-Poisson and for the estimates based on the thinned MCMC sample (right panel). The bias of PS-LOO-CV appears to be more pronounced than that of the original CPO approach (LOO-CV). The wAIC estimate is generally intermediate,

| | | | LOO-CV | | | | | | wAIC | | | | |
|---|---|---|---|---|---|---|---|---|---|---|---|---|---|
| | n | T | est[1] | bias[2] | stdev[3] | ESS[4] | %ESS< 10[5] | %$\hat{\kappa} > 0.7$[6] | est[1] | bias[2] | stdev[3] | ESS[4] | %ESS< 10[5] |
| LG | 500 | 100 | -24.128 | 0.0018 | 0.0034 | 81.5 | 0.00 | 0.10 | -24.126 | -0.0015 | 0.0033 | 83.2 | 0.00 |
| | 500 | 1000 | -24.128 | 0.0004 | 0.0013 | 806.4 | 0.00 | 0.00 | -24.127 | -0.0003 | 0.0012 | 826.9 | 0.00 |
| | 1000 | 100 | -24.783 | 0.0008 | 0.0022 | 89.1 | 0.00 | 0.00 | -24.782 | -0.0007 | 0.0022 | 89.8 | 0.00 |
| | 1000 | 1000 | -24.782 | 0.0002 | 0.0007 | 888.0 | 0.00 | 0.00 | -24.782 | -0.0001 | 0.0007 | 896.6 | 0.00 |
| | 4000 | 100 | -24.841 | 0.0002 | 0.0003 | 96.9 | 0.00 | 0.01 | -24.840 | -0.0002 | 0.0002 | 97.0 | 0.00 |
| | 4000 | 1000 | -24.840 | 0.0000 | 0.0001 | 968.2 | 0.00 | 0.00 | -24.840 | -0.0000 | 0.0001 | 969.2 | 0.00 |
| | 16000 | 100 | -24.697 | 0.0000 | 0.0001 | 99.2 | 0.00 | 0.00 | -24.697 | -0.0000 | 0.0001 | 99.2 | 0.00 |
| | 16000 | 1000 | -24.697 | 0.0000 | 0.0000 | 991.9 | 0.00 | 0.00 | -24.697 | -0.0000 | 0.0000 | 992.0 | 0.00 |
| GTR | 500 | 100 | -24.111 | 0.0107 | 0.0030 | 68.2 | 4.14 | 3.35 | -24.102 | -0.0052 | 0.0023 | 72.1 | 1.26 |
| | 500 | 1000 | -24.111 | 0.0024 | 0.0025 | 658.2 | 0.36 | 2.00 | -24.105 | -0.0011 | 0.0025 | 707.3 | 0.00 |
| | 1000 | 100 | -24.726 | 0.0036 | 0.0044 | 77.3 | 0.43 | 0.62 | -24.722 | -0.0022 | 0.0036 | 79.9 | 0.14 |
| | 1000 | 1000 | -24.727 | 0.0012 | 0.0017 | 760.5 | 0.00 | 0.25 | -24.724 | -0.0006 | 0.0010 | 791.2 | 0.00 |
| | 4000 | 100 | -24.736 | 0.0008 | 0.0008 | 91.4 | 0.03 | 0.07 | -24.734 | -0.0006 | 0.0008 | 92.1 | 0.01 |
| | 4000 | 1000 | -24.735 | 0.0002 | 0.0004 | 912.1 | 0.00 | 0.02 | -24.735 | -0.0002 | 0.0005 | 919.3 | 0.00 |
| | 16000 | 100 | -24.573 | 0.0002 | 0.0001 | 97.5 | 0.00 | 0.00 | -24.573 | -0.0001 | 0.0001 | 97.6 | 0.00 |
| | 16000 | 1000 | -24.573 | 0.0001 | 0.0001 | 974.5 | 0.00 | 0.00 | -24.573 | -0.0001 | 0.0001 | 975.5 | 0.00 |
| CAT | 500 | 100 | -23.832 | 0.0604 | 0.0219 | 60.2 | 27.85 | 12.35 | -23.810 | -0.0125 | 0.0172 | 63.6 | 8.61 |
| | 500 | 1000 | -23.833 | 0.0195 | 0.0115 | 574.8 | 6.99 | 11.80 | -23.816 | -0.0014 | 0.0028 | 619.6 | 0.00 |
| | 1000 | 100 | -24.385 | 0.0244 | 0.0084 | 66.5 | 18.01 | 8.05 | -24.396 | -0.0100 | 0.0082 | 68.3 | 6.56 |
| | 1000 | 1000 | -24.400 | 0.0076 | 0.0041 | 644.0 | 3.78 | 7.58 | -24.408 | -0.0013 | 0.0035 | 667.7 | 0.00 |
| | 4000 | 100 | -24.274 | 0.0112 | 0.0034 | 77.2 | 6.80 | 3.10 | -24.281 | -0.0060 | 0.0023 | 77.1 | 4.31 |
| | 4000 | 1000 | -24.274 | 0.0027 | 0.0018 | 759.8 | 1.00 | 2.47 | -24.284 | -0.0010 | 0.0022 | 759.7 | 0.03 |
| | 16000 | 100 | -24.000 | 0.0037 | 0.0009 | 85.5 | 2.24 | 1.03 | -24.004 | -0.0036 | 0.0007 | 84.4 | 2.48 |
| | 16000 | 1000 | -24.001 | 0.0009 | 0.0004 | 848.1 | 0.16 | 0.70 | -24.006 | -0.0007 | 0.0005 | 835.4 | 0.01 |

[1]: estimated; [2]: estimated Monte Carlo bias; [3]: estimated Monte Carlo standard deviation; [4]: mean effective sample size (ESS) across sites; [5]: percentage of sites with ESS $< 10$; [6]: percentage of sites with $\hat{\kappa} > 0.7$.

Table 1: Numerical estimates and Monte Carlo statistics for the LOO-CV and wAIC scores, for the scaling experiment on the Metazoa dataset.

lower than LOO-CV and higher than 2-step CV. This suggests that the asymptotic error of the wAIC is small compared to the numerical biases of the non-asymptotic approaches. In the case of GTR, there is no clear trend, presumably because the Monte Carlo estimation bias, which is much smaller than in the case of CAT-Poisson, is dominated by sampling variance (i.e. variance over data replicates).

Supplementary Figure 2 shows the fit of LOO-CV, the wAIC and 2-step-CV separately for each replicate, for the full MCMC sample ($T = 1000$) against the thinned sample ($T = 100$). Plotting the two settings against each other allows for an evaluation of the stability of the estimators upon thinning, and gives also a visual representation of the bias: the bias is expected to decrease linearly as a function of $T$, and thus, a negatively (resp. positively) biased estimator is expected to lean toward the upper left (resp. bottom right) side of the first diagonal. For GTR, all estimators appear to be very stable. For the numerically more challenging CAT-Poisson model, the wAIC appears to be the most stable estimator, falling along the diagonal, followed by 2-step CV (whose negative bias is visible), then LOO-CV (showing a positive bias). In all cases, de-biasing the estimate appears to be useful (bottom-right panel, CAT-Poisson), or at least not harmful when not necessary (bottom left, GTR).

Alternatively, Supplementary Figures 3 and 4 show the fit for each replicate for the wAIC against LOO-CV (Supplementary Figure 3) or for 2-step CV against LOO-CV (Supplementary Figure 4). This confirms that wAIC and LOO-CV are very similar to each other, once de-biased (Supplementary Figures 3) – except for two points corresponding to low sample size ($n = 500$), suggesting that the numerical instabilities about which low ESS or high $\kappa$ are warning about (Supplementary Table 1) may occasionally have a non-negligible impact in this regime of data size and for complex models. The comparison of LOO-CV with 2-step CV separately for each replicate (Supplementary Figure 4) is more noisy, owing to the additional sampling variance contributed by the training on an independent dataset for 2-step CV. Of note, this noise gives an indirect measure of the sampling variance, which in the present case is larger than the occasional numerical instabilities mentioned above for $n = 500$ (compare with Supplementary Figure 3). Conversely, this additional variance decreases with data size, such that, for larger datasets ($n \geq 4000$ – corresponding to the points located in the upper right of each quadrant), the estimates again appear to match closely between the two methods, once de-biased.

Altogether, once the estimators have been de-biased, both the LOO-CV and the wAIC estimates are reasonably accurate and robust to varying levels of thinning. For datasets larger than 4000

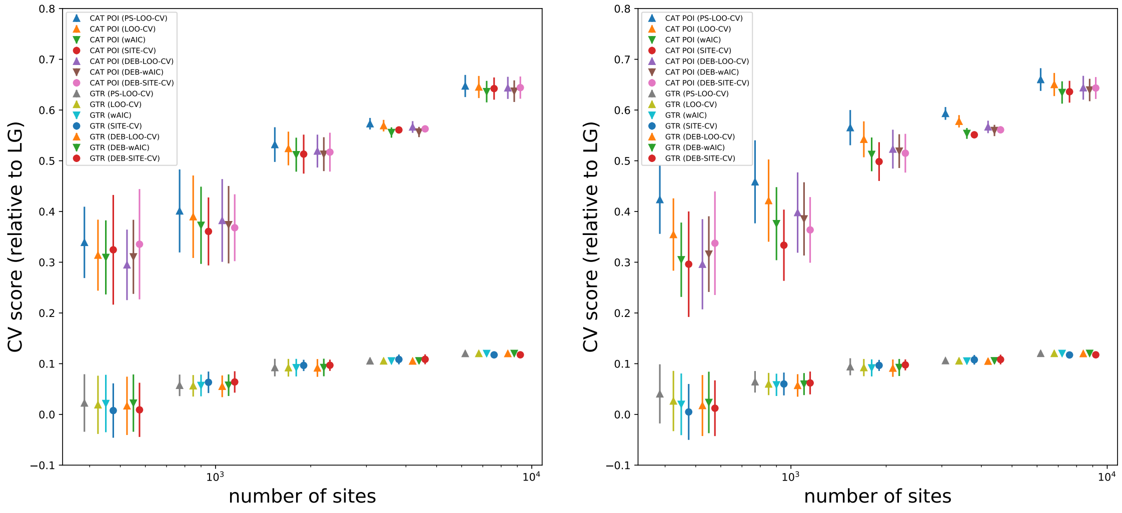


Figure 1: PS-LOO-CV, LOO-CV, wAIC and 2-step CV estimates for the GTR and CAT-Poisson models (relative to LG), using $T = 1000$ samples, i.e. without thinning (left panel) or $T = 100$ samples, i.e. with 10-fold thinning (right panel), as a function of datasize. For each data size, the 7 estimates are, from left to right: PS-LOO-CV, raw LOO-CV, wAIC, 2-step CV, then de-biased LOO-CV, wAIC and 2-step CV.

aligned positions, they seem to be virtually unaffected by thinning and identical to each other and to the independent 2-step approach. For smaller data size, ($n < 1000$), there seems to be some instances of numerical instabilities – which remain small in size, however, compared to the sampling variance. The wAIC and 2-step CV appear to be more stable than LOO-CV and PS-LOO-CV. PS-LOO-CV does not seem to bring an improvement over LOO-CV in the present case, as it shows a stronger positive bias than LOO-CV. On the other hand, through the estimated $\hat{\kappa}$, it may offer a useful additional quality check for the stability of the estimation. Finally, the asymptotic error entailed by the wAIC appears to be negligible for $n > 5000$.

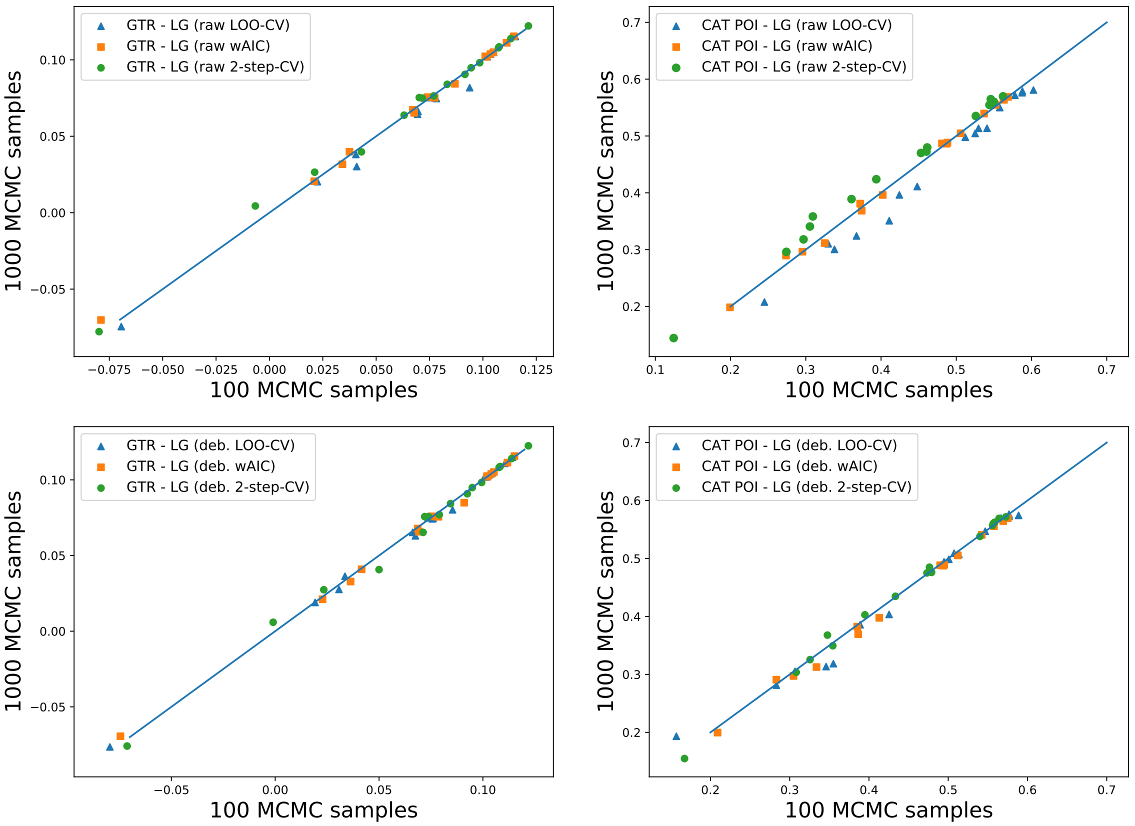


Figure 2: raw (top) and de-biased (bottom) LOO-CV, wAIC and 2-step CV estimates, for the GTR (left) and CAT-Poisson (right) models (relative to LG), using $T = 1000$ samples (y-axis) versus $T = 100$ samples (x-axis). Of note, all estimates falling in the upper right quadrant of each panel (i.e. above $x = 0.5$ and $y = 0.5$) correspond to datasets larger than 4000 aligned positions.

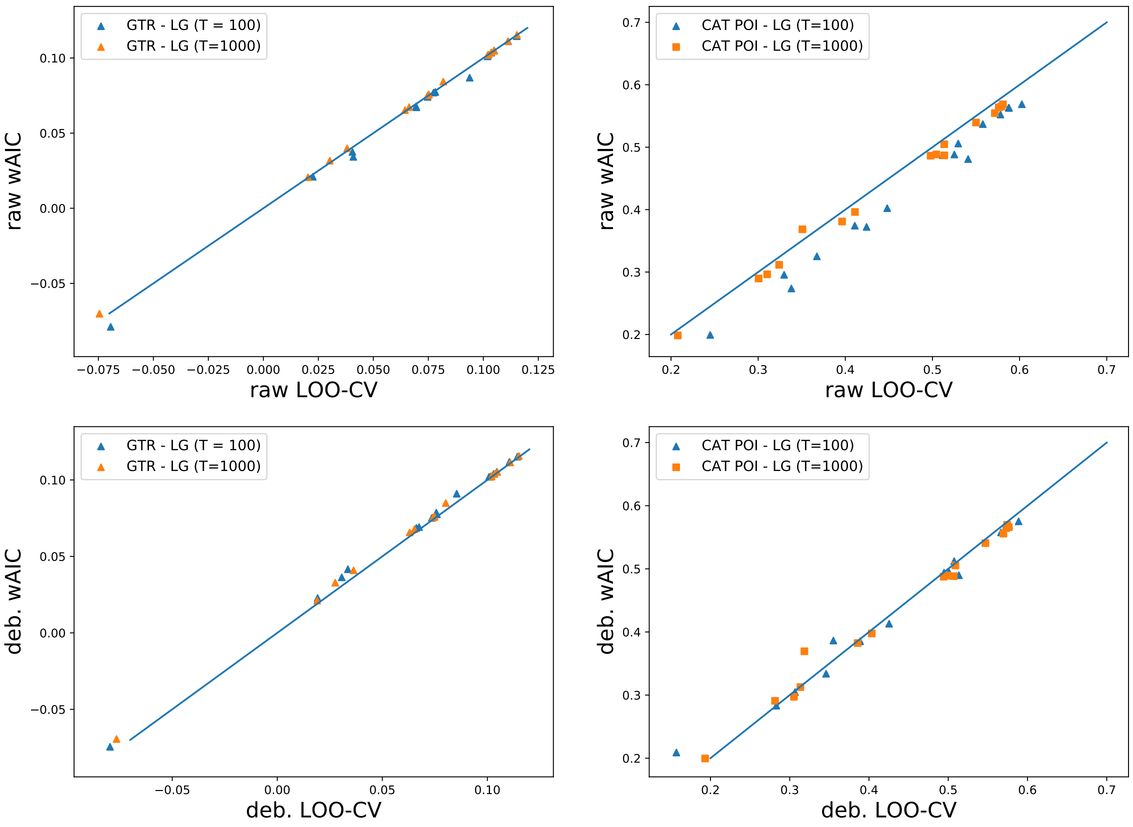


Figure 3: raw (top) and de-biased (bottom) estimates of the wAIC (y-axis) against LOO-CV (x-axis) for the GTR (left) and CAT-Poisson (right) models (relative to LG), using $T = 1000$ samples or $T = 100$ samples. Of note, all estimates falling in the upper right quadrant of each panel (i.e. above $x = 0.5$ and $y = 0.5$) correspond to datasets larger than 4000 aligned positions.

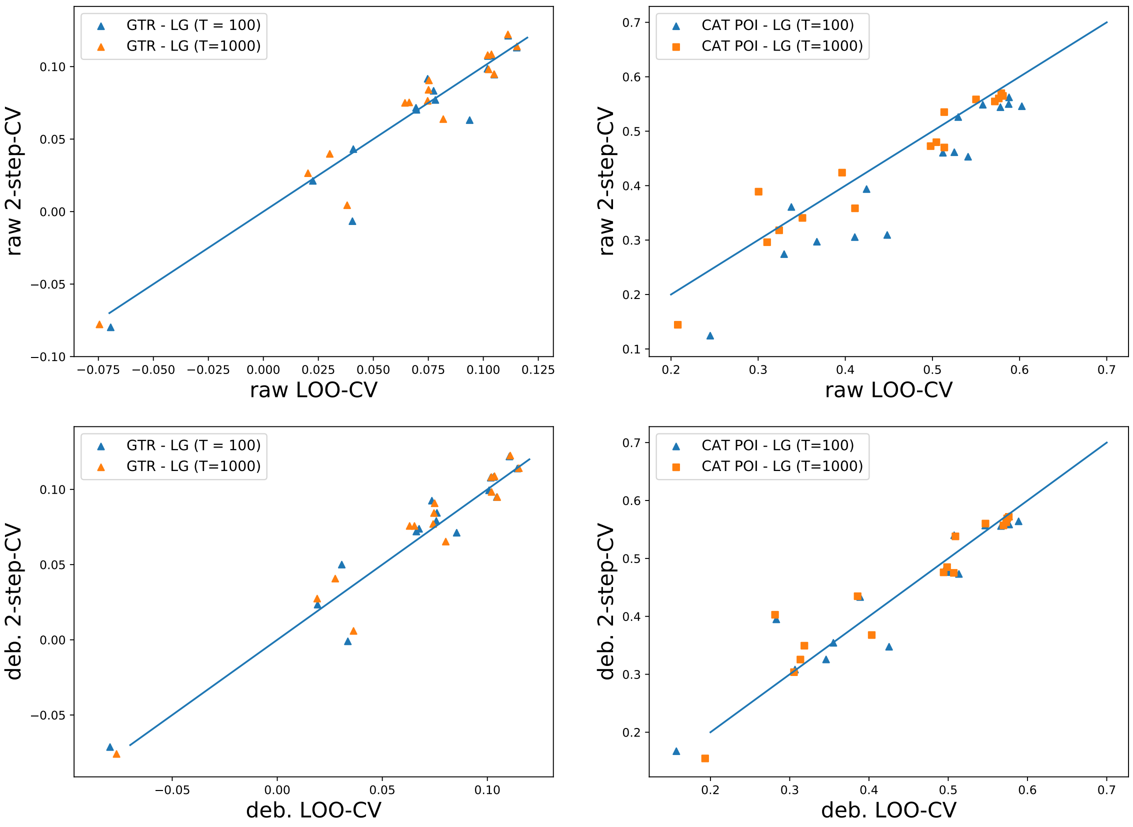


Figure 4: raw (top) and de-biased (bottom) estimates of 2-step-CV (y-axis) against LOO-CV (x-axis) for the GTR (left) and CAT-Poisson (right) models (relative to LG), using $T = 1000$ samples or $T = 100$ samples. Of note, all estimates falling in the upper right quadrant of each panel (i.e. above $x = 0.5$ and $y = 0.5$) correspond to datasets larger than 4000 aligned positions.

#### 4 Additional theoretical results on the expected predictive fit

As in the main document, the general formula for the expected predictive fit on a dataset of size $r$ upon training on an independent dataset of size $q$ is noted:

$$C(q, r) = \frac{1}{r} E\left[\ln p(Y_r \mid Y_q)\right] \tag{7}$$

where $Y_q$ and $Y_r$ are two independent datasets randomly drawn from the population, of size $q$ and $r$, respectively.

The predictive fit follows a mathematical identity that can be derived as follows. Consider a dataset made of $n = q + r$ independent data points, $X_{1:n}$ and assume for notational convenience that the first $q$ data points were used for training, and the last $r$ for validation. By the chain rule, the joint probability of the validation set can be expressed in terms of a sequential product of the marginal likelihoods of each of individual observations.

$$p(X_{q+1:n} \mid X_{1:r}) = \prod_{i=q+1}^{n} p(X_i \mid X_{1:i-1}) \tag{8}$$

or, on a logarithmic scale and on a per-site basis:

$$\frac{1}{r} \ln p(X_{q+1:n} \mid X_{1:r}) = \frac{1}{r} \sum_{i=q+1}^{n} \ln p(X_i \mid X_{1:i-1}) \tag{9}$$

Taking the expectation over $X$ gives:

$$C(q, r) = \frac{1}{r} \sum_{i=q+1}^{n} C(i-1, 1). \tag{10}$$

Thus, when training on a dataset of size $q$ and then validating on a dataset of size $r$, everything happens as if one were progressively training the model over a series of data sets of increasing size, between $q$ and $q+r=n$, each time computing the fit on the next individual data point and, finally, averaging over the whole series. Given that $C(q, 1)$ is monotonously increasing as a function of $q$, this implies in particular that $C(q, r) \geq C(q, 1)$ for any $r$, and more generally, that $C(q, s) \geq C(q, r)$ for any $r$ and $s$ such that $s \geq r$.

This property, which is a direct consequence of the chain rule, explains why joint k-fold CV advertises a higher apparent fit than site-wise k-fold CV, and this, in spite of the fact that training was done on a dataset of identical size in both cases. Another way to say it is that k-fold CV behaves mathematically as if some cryptic training were also occurring on the validation dataset. This behavior of k-fold CV may be considered problematic on purely conceptual grounds, as it violates the principle of a clear the separation between training and validation, which was the whole point of cross-validation. Sitewise k-fold CV does not have this problem.

For the same reason, the chain rule on $C(q, r)$ also provides a justification for defining the Bayes generalization error as the posterior mean KL-divergence upon seing a single new data point, i.e. $B_g = H^* - C(n, 1)$, such as proposed by Watanabe (2010). Consider instead the alternative definition of $B_g$ as the posterior mean KL-divergence upon seeing a new data set of same size as the current one, i.e. $B_g' = H^* - \frac{1}{n} C(n, n)$. By the chain rule, however, this would amount to training the model over a data set of size larger than $n$.

### 5 Analytical results under the normal model

The normal model assumes that the $n$ observations $(X_i)_{i=1..n}$ are i.i.d. normal of known variance:

$$X_i \sim N(\theta, \sigma^2 I_p),$$

where $\theta$ is fixed and equal to 0 under model $M_1$ and is unknown and endowed with a normal prior of covariance $\Sigma_0 = \delta^2 I_p$ under model $M_2$:

$$\theta \sim N(0, \delta^2 I_p)$$

In the following, the analytical results under the normal model are first derived in the univariate case. Thus, the random variable $X_i$ and its observed value $x_i$, for $i = 1..n$, are then real numbers. The generalization to the multivariate case is straighforward, since the model is spherical.

For notational convenience, the analytical results are expressed in terms of the precision parameters $\tau = 1/\sigma^2$ and $\omega = 1/\delta^2$. For any real function $f$, the empirical mean over the $n$ observed values is noted:

$$\langle f(x) \rangle_n = \frac{1}{n} \sum_i f(x_i)$$

Model $M_1$ does not have free parameters. The marginal likelihood under this model is thus simply:

$$p(X_{1:n} \mid M_1) = \left( \frac{\tau}{2\pi} \right)^{\frac{n}{2}} e^{-\frac{n\tau}{2} \langle x^2 \rangle_n}$$

or, on a logarithmic scale:

$$\ln p(X_{1:n} \mid M_1) = \frac{n}{2} \ln \frac{\tau}{2\pi} - \frac{n\tau}{2} \langle x^2 \rangle_n$$

The likelihood under model $M_2$ is:

$$p(X_{1:n} \mid \theta, M_2) \;=\; \left(\frac{\tau}{2\pi}\right)^{\frac{n}{2}} e^{-\frac{n\tau}{2}\langle (x-\theta)^2\rangle_n}$$

the prior:

$$p(\theta \mid M_2) \;=\; \left(\frac{\omega}{2\pi}\right)^{\frac{1}{2}} e^{-\frac{\omega}{2}\theta^2}$$

and thus the marginal likelihood:

$$\begin{aligned} p(X_{1:n} \mid M_2) &= \int p(X_{1:n} \mid \theta, M_2) p(\theta \mid M_2) d\theta \\ &= \left(\frac{\tau}{2\pi}\right)^{\frac{n}{2}} \left(\frac{\omega}{2\pi}\right)^{\frac{1}{2}} \int e^{-\frac{1}{2}\left(n\tau\langle (x-\theta)^2\rangle_n + \omega\theta^2\right)} d\theta \end{aligned}$$

Relying on the identity:

$$\omega\theta^2 + n\tau\langle (x-\theta)^2\rangle_n \;=\; (\omega + n\tau)\left[(\theta - \tilde{x}_n)^2 + \tilde{v}_n\right]$$

where

$$\tilde{x}_n \;=\; \frac{n\tau}{\omega + n\tau}\langle x\rangle_n \tag{11}$$

$$\tilde{v}_n \;=\; \frac{n\tau}{\omega + n\tau}\langle x^2\rangle_n - \left(\frac{n\tau}{\omega + n\tau}\right)^2 \langle x\rangle_n^2 \tag{12}$$

and integrating out the normal component that depends on $\theta$ gives:

$$p(X_{1:n} \mid M_2) \;=\; \left(\frac{\tau}{2\pi}\right)^{\frac{n}{2}} \left(\frac{\omega}{2\pi}\right)^{\frac{1}{2}} \int e^{-\frac{1}{2}(\omega+n\tau)\left[(\theta-\tilde{x}_n)^2 + \tilde{v}_n\right]} d\theta \tag{13}$$

$$\;=\; \left(\frac{\tau}{2\pi}\right)^{\frac{n}{2}} \left(\frac{\omega}{2\pi}\right)^{\frac{1}{2}} \left(\frac{2\pi}{\omega + n\tau}\right)^{\frac{1}{2}} e^{-\frac{1}{2}(\omega+n\tau)\tilde{v}_n} \tag{14}$$

or, on the logarithmic scale:

$$\ln p(X_{1:n} \mid M_2) \;=\; \frac{n}{2}\ln\frac{\tau}{2\pi} + \frac{1}{2}\ln\frac{\omega}{\omega + n\tau} - \frac{\omega + n\tau}{2}\tilde{v}_n$$

The log of the Bayes factor is thus equal to:

$$\ln BF \;=\; \frac{1}{2}\ln\frac{\omega}{\omega + n\tau} + \frac{n\tau}{2}\langle x^2\rangle_n - \frac{\omega + n\tau}{2}\tilde{v}_n \tag{15}$$

$$\;=\; \frac{1}{2}\left(\ln\frac{\omega}{\omega + n\tau} + \frac{(n\tau)^2}{\omega + n\tau}\langle x\rangle_n^2\right) \tag{16}$$

where the definition of $v_n$ above was use to simplify out the term in $\langle x^2\rangle_n$.

For the cross-validation score, assuming that the data have been re-ordered, such that the first $q$ observations correspond to the training set $X^t$ and the last $r$ to the validation set $X^v$, with $q + r = n$, then the cross-validation log likelihood score for a given arbitrary model is simply:

$$\ln p(X^v \mid X^t) \;=\; \ln p(X_{1:n}) - \ln p(X_{1:q})$$

Equivalently, when comparing two models:

$$\begin{aligned}\Delta \ln p(X^v \mid X^t) &= \Delta \ln p(X_{1:n}) - \Delta \ln p(X_{1:q}) \\ &= \ln BF_{1:n} - \ln BF_{1:q}\end{aligned}$$

The score can thus be computed directly, simply by invoking equation 16 successively for each of the two terms.

From equation 13, and in particular from the dependence in $\theta$ under the integral sign, it can also be seen that the posterior distribution is normal, of mean $\tilde{x}_n$ and variance $(\omega + n\tau)^{-1}$. More generally, the partial posterior obtained by conditioning on a subset of the data $X_{1:q}$ is also normal:

$$\theta \mid X_{1:q} \sim \texttt{Normal}\left(\tilde{x}_q, (\omega + q\tau)^{-1}\right), \tag{17}$$

where $\tilde{x}_q$ is obtained by replacing $n$ by $q$ in equation 11. This result is used in the MCMC sampling method for the normal model (see Materials and Methods section).

The analytical results presented thus far give the logarithm of the marginal likelihood and cross-validation scores for a given dataset $X$. The results presented on Figure 1 of main manuscript rely on the expectations of these quantities, upon resampling $X$ from the population. Since $\langle x \rangle_n$ is the empirical mean of $n$ random variables of mean $\theta$ and variance $\sigma^2$, it has itself mean $\theta$ and variance $\frac{\sigma^2}{n}$ and thus its square has expectation:

$$E\left[\langle x \rangle_n^2\right] = \theta^2 + \frac{\sigma^2}{n}$$

or equivalently:

$$n\tau E\left[\langle x \rangle_n^2\right] = n\tau\theta^2 + 1$$

Plugging this identity into equation 16 gives the expected log Bayes factor:

$$E[\ln BF] = \frac{1}{2}\left(\ln \frac{\omega}{\omega + n\tau} + \frac{n\tau}{\omega + n\tau}(n\tau\theta^2 + 1)\right)$$

On a per-site basis, and using the notation introduced in the main text:

$$C(0,n) = E[\Delta m] = \frac{1}{n} E[\ln BF] \tag{18}$$

Finally, the expected cross-validation score is defined as:

$$C(q,r) = \frac{1}{r} E\left[\Delta \ln p(X^v \mid X^t)\right] = \frac{1}{r}\left(E[\Delta \ln p(X_{1:q+r})] - E[\Delta \ln p(X_{1:q})]\right) \tag{19}$$

$$= \frac{1}{r}\left[(q+r)C(0,q+r) - qC(0,q)\right] \tag{20}$$

which, together with equation 18, gives the expected score for all cross-validation schemes. These were not further simplified.

### References

Vehtari, A., Gelman, A. & Gabry, J. 2016 Practical Bayesian model evaluation using leave-one-out cross-validation and WAIC. *Stat Comput*, **27**(5), 1413–1432.
